# A complex regulatory network governs the production of an antibiotic with unusual cell-density-dependence

**DOI:** 10.1101/2023.12.13.571536

**Authors:** Hindra, Marie A. Elliot

## Abstract

*Streptomyces* bacteria are renowned both for their antibiotic production capabilities, and for their cryptic metabolic potential. Here, we leveraged the activity of an Lsr2 knockdown construct to enhance antibiotic production in the wild *Streptomyces* isolate WAC07094. We determined the new activity stemmed from increased levels of the angucycline-like family member saquayamycin. Saquayamycin has both antibiotic and anti-cancer activities, and intriguingly, beyond Lsr2-mediated repression, we found saquayamycin production was also suppressed at high density on solid or in liquid growth media. This density-dependent control was exerted at the level of the cluster-situated regulatory gene *sqnR* and was mediated in part through the activity of the PhoRP two component regulatory system; deleting *phoRP* led to both constitutive antibiotic production and *sqnR* expression, suggesting that PhoP functions to repress the expression of *sqnR* at high cell density. We further discovered that magnesium supplementation could also alleviate this cell density dependence, although its action was independent of PhoP. Finally, we revealed that the nitrogen-responsive regulators GlnR and AfsQ1 could relieve the repression exerted by Lsr2 and PhoP. This unusual density-dependent production of saquayamycin was not unique to WAC07094; we found that saquayamycin production by another wild isolate was also density-dependent, suggesting this spatial control may serve an important ecological function in their native environments.

**IMPORTANCE:** *Streptomyces* specialized metabolic gene clusters are subject to complex regulation, and their products are frequently not observed under standard laboratory growth conditions. For the wild *Streptomyces* isolate WAC07094, production of the angucycline-family compound saquayamycin is subject to a unique constellation of control factors. Notably, it is produced primarily at low cell density, in contrast to the high cell density production typical of most antibiotics. This unusual density dependence is conserved in other saquayamycin producers and is driven by the pathway-specific regulator SqnR, whose expression is influenced by both nutritional and genetic elements. Collectively, this work provides new insights into an intricate regulatory system governing antibiotic production and indicates there may be benefits to including low density cultures in antibiotic screening platforms.

## INTRODUCTION

Microbial specialized metabolites are important sources and inspirations for antibiotic discovery and development (1–3). *Streptomyces* bacteria, and their actinobacterial relatives, are major contributors to the antibiotics used in clinical and agricultural settings. These microbes also produce a wide array of biologically active molecules having antifungal, anticancer, immunosuppressant and herbicide utility. The synthesis of any specialized metabolite is a complex and highly regulated process, and the production of these compounds is thought to confer their producers with a competitive advantage in their natural environment. Consequently, these molecules are often not produced at significant levels under conventional laboratory growth conditions.

Antibiotic production, and specialized metabolism more broadly, are integrated into the *Streptomyces* developmental cycle, where they are temporally linked with the onset of aerial (reproductive) growth (4). The transition from vegetative hyphal growth to the raising of aerial hyphae, formation of reproductive spores, and initiation of specialized metabolism, is in turn linked to conditions of nutrient depletion, including low levels of carbon, nitrogen, and phosphate (5–7). Quorum-sensing has also been tied to both development and antibiotic control (8), implicating a role for cell signalling and cell density in regulating specialized metabolism. Most characterized specialized metabolite biosynthetic clusters are subject to multi-level regulation (9). These clusters often encode dedicated cluster-situated regulators, many of which are transcriptional activators. These regulatory genes in turn, are subject to control by multiple globally acting transcription factors, ranging from nutrient-sensing regulators like DasR (*N*-acetylglucosamine) (7), GlnR (nitrogen) (10) and PhoRP (phosphate) (11), through to nucleoid-associated proteins like Lsr2 (12). There are further instances when binding sites for these different regulators overlap (13–14); this adds to the regulatory complexity governing specialized metabolism and provides opportunities for both distinct regulatory inputs, and interplay between different regulators.

The nucleoid-associated protein Lsr2 is conserved throughout the actinobacteria and is functionally similar to the well-studied H-NS protein from *Escherichia coli* in serving as a silencer of horizontally acquired genes (15). In the streptomycetes, Lsr2 also represses the expression of many genes involved in specialized metabolism (12). This repression can be overcome by sequestering the native protein through overexpression of a dominant negative variant that is dimerization/oligomerization-proficient but DNA-binding defective (12). Constitutively expressing this ‘Lsr2 knockdown’ construct has proven useful in stimulating the production of new metabolites by wild *Streptomyces* species, presumably by relieving the biosynthetic cluster silencing exerted by the native protein (12).

We introduced this Lsr2 knockdown construct into a suite of wild *Streptomyces* isolates and identified one recipient that had enhanced antibiotic production relative to its parent strain. Unusually, its antibiotic production was most pronounced when cells were growing at low density; there was no activity seen for high-density cells, which is the converse of what is usually seen for antibiotic synthesis. We first determined that the antibiotic with these intriguing production characteristics was saquayamycin, a molecule that belongs to the angucycline family of natural products (16–17). To better understand the mechanistic basis for the low cell-density production of saquayamycin, we probed the effects of both environmental factors (diverse nutrients), and genetic regulators. We found that magnesium salts relieved the density dependence, and significantly enhanced saquayamycin production yields. From a regulatory perspective, beyond the stimulatory effects observed for Lsr2 knockdown, we further identified the two-component system PhoP/PhoR as a key regulator of saquayamycin. Notably, we determined that the low cell-density dependence of saquayamycin production was conserved in other saquayamycin producers and was subject to the same regulatory and nutritional controls in these other species.

## RESULTS

### Lsr2 knockdown promotes new antibiotic production for WAC07094 on agar medium

In an effort to activate silent biosynthetic gene clusters, we introduced our Lsr2 knockdown plasmid into a suite of wild *Streptomyces* isolates and screened these genetically modified strains for new antibiotic production. We found that introducing this construct into *Streptomyces* WAC07094 (WAC: Wright Actinomycete Collection) led to new pigmentation and strong antibiotic activity against Gram-positive bacteria, including methicillin-resistant *Staphylococcus aureus* and vancomycin-resistant Enterococci, in contrast to its plasmid-free or empty plasmid-containing parental strain which were unpigmented and had either no or significantly reduced levels of growth-inhibition (**Figure 1A; Figure S1**).

**Figure 1.**
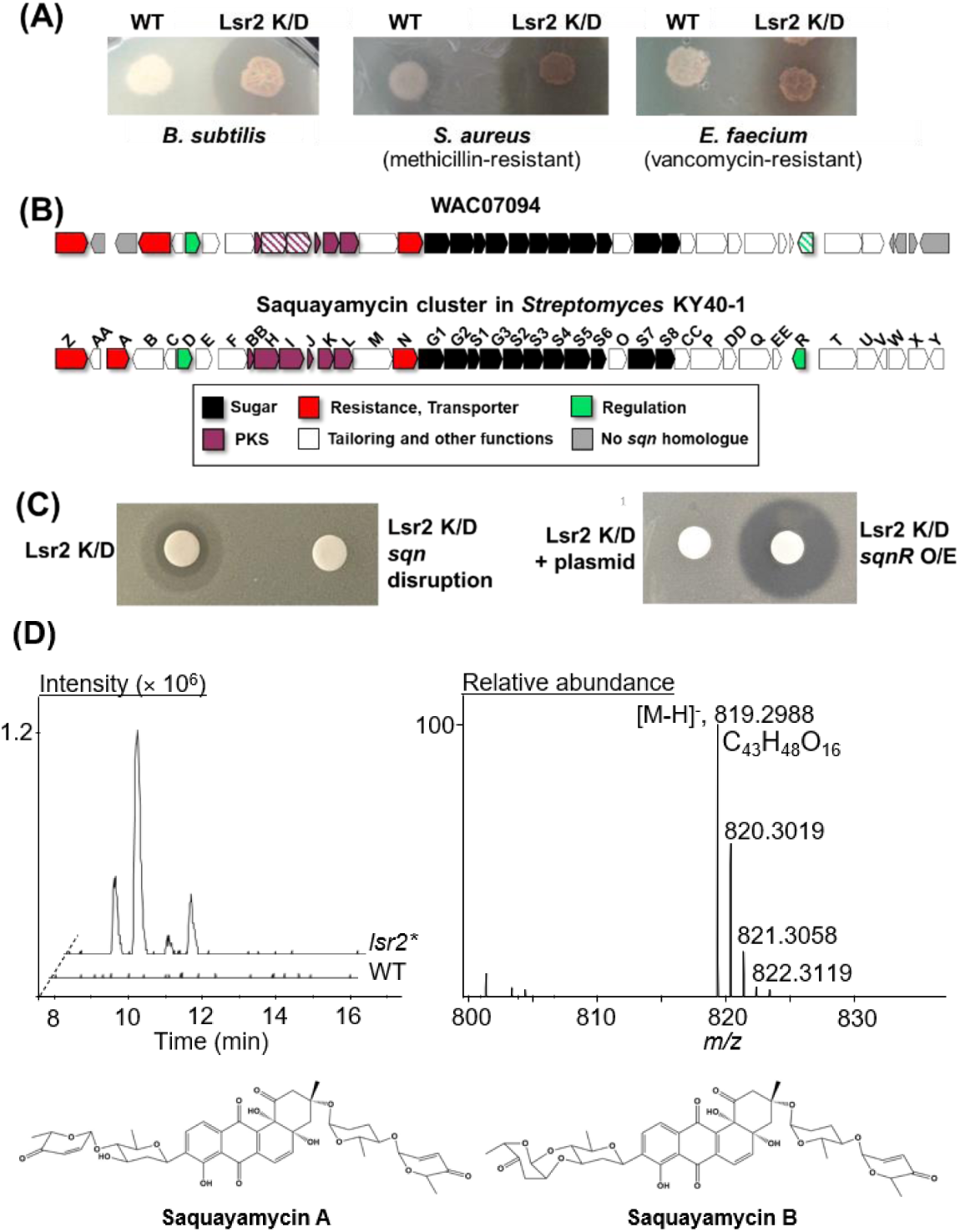
Saquayamycin production by *Streptomyces* sp. WAC07094. **(A)** Antibiotic bioassay in which wild type (WT) WAC7094 and Lsr2 knockdown (K/D) strains (overexpressing a dominant negative variant of *lsr2*) were spotted to Bennett’s medium, grown for 3 days, and then overlaid with the sensitive indicator strains *Bacillus subtilis* (left), methicillin-resistant *Staphylococcus aureus* (middle) or vancomycin-resistant *Enterococcus faecium* (right). **(B)** Schematic diagram of the saquayamycin biosynthetic cluster in *Streptomyces* KY40-1 (bottom), compared with the equivalent cluster from WAC07094 (top). Genes from the WAC07094 strain studied here are indicated with hashed lines. **(C)** Antibiotic bioassay using *B. subtilis* as the indicator strain, together with crude extracts spotted to filter discs, from the *sqnHI* disruption mutant and the *sqnR* overexpression strains (both carrying the Lsr2 knockdown construct) grown for 3 days on Bennett’s medium, compared with their relevant controls. **(D)** LC-MS analysis of crude extracts prepared from WAC07094 grown for 3 days on Bennett’s agar (agar + biomass). Left: chromatograms of [M-H]^-^ ions extracted at *m/z* 819. Right: Mass spectra of the *m/z* 819 peak and its assigned chemical formula. Bottom: structures of saquayamycin A and B.

**Figure S1.**
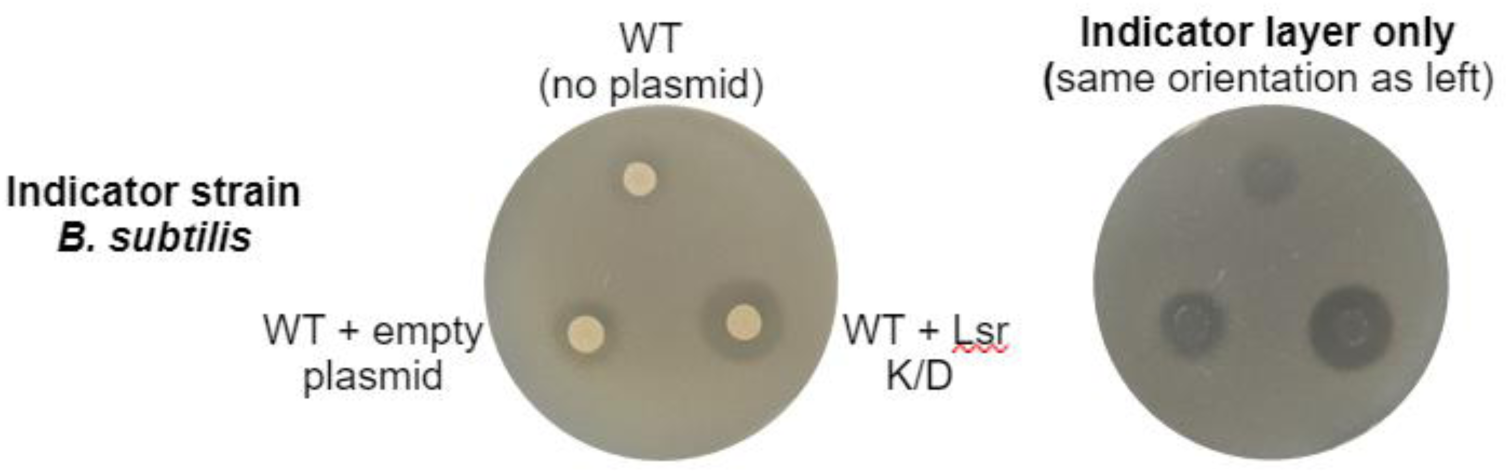
Antibiotic bioassay of *Streptomyces* WAC07094 strains. Antibiotic activity of wild type (WT) WAC07094 (top) was compared with that of an empty plasmid-carrying variant (left) and one carrying an Lsr2 knockdown (K/D)-expressing construct (right) using a ‘sandwich-based’ bioassay, where equal numbers of spores for each strain were spotted to Bennett’s medium, grown for 3 days, and then overlaid with nutrient agar infused with the indicator strain *Bacillus subtilis* (left image). The image to the right shows only the *B. subtilis*-infused nutrient agar (indicator layer) after being separated from the ’sandwich’ to better illustrate the zones of inhibition.

To identify the specialized metabolite (and its associated biosynthetic gene cluster) responsible for this new bioactivity, the WAC07094 genome was sequenced. The final genome assembly consisted of 4 contigs having a total length of 9,599,672 bp (accession no. JAVMJZ000000000), including 18 biosynthetic gene clusters predicted using antiSMASH 5.0 (18) (**Supplementary Table 1**). In analyzing the annotated genome, we found that WAC07094 was unusual in encoding three *lsr2*-like genes, with two sharing strong similarity with the *lsr2* and *lsrL* genes encoded by other streptomycetes. The third gene encoded a protein that was more similar to Lsr2 than LsrL, and was dubbed ’LsrS’, for <u>Lsr</u>2 <u>s</u>imilar (**Figure S2**). The promoter region of *lsrS* was highly similar (95% identity over 240 nt with no gaps) to that of the *lsr2* promoter, but the flanking genes differed from those adjacent to the canonical *lsr2* and included a putative transposase-encoding gene, suggesting that *lsrS* may have arisen through a genome duplication and transposition event.

**Figure S2.**
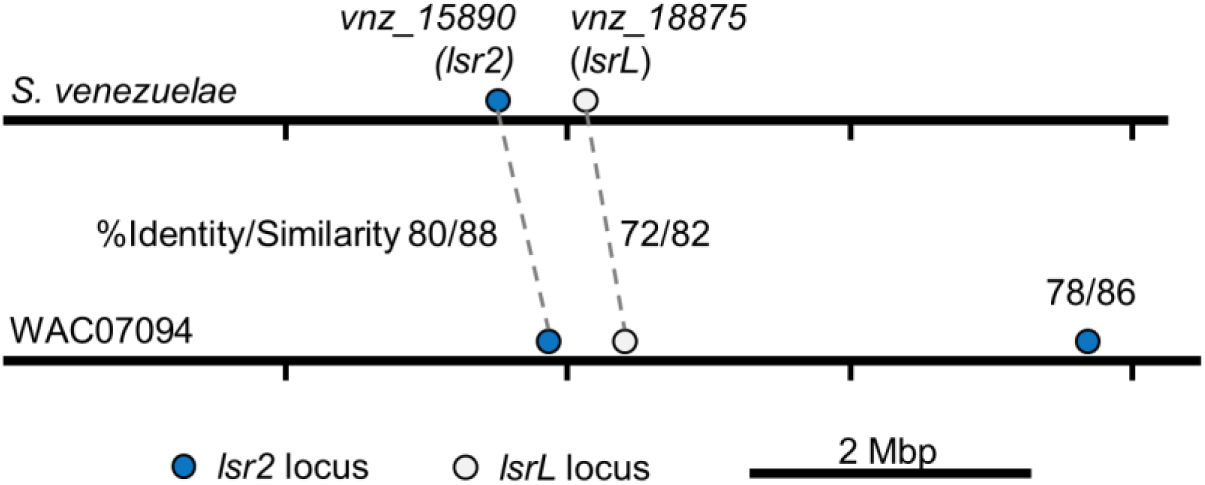
Relative locations and similarities of Lsr2 and its homologues in *S. venezuelae* and WAC07094. Schematic diagram illustrating the relative position of *lsr2* and *lsrL* homologues in the chromosomes of *S. venezuelae* and WAC07094. For WAC07094, the percent identity/similarity are indicated, relative to their *S. venezuelae* equivalents. For LsrS (the third Lsr2-like protein encoded by WAC07094), the identity/similarity values are relative to *S. venezuelae* Lsr2.

We knew from previous work that Lsr2 binds AT-rich regions (12). Consequently, we assessed the relative AT content associated with each of the 18 predicted biosynthetic clusters and found that several clusters had AT-rich regions that could be targeted by Lsr2 (**Supplementary Table 1**). As Lsr2 typically represses gene expression, we predicted that Lsr2 knockdown would be associated with increased expression of at least one of these biosynthetic clusters, leading to the observed antibiotic activity. To identify this cluster, RNA was isolated from wild type and Lsr2 knockdown strains after three days (when antibacterial activity was observed), and these samples were subjected to RNA sequencing. Comparing the levels of sequencing reads (Accession no. PRJNA1009436) between strains revealed that a polyketide synthase-containing gene cluster with an extended AT-rich region, was upregulated in the Lsr2 knockdown strain (**Figure S3**). This cluster bore 85% similarity to the previously characterized saquayamycin biosynthetic gene cluster (MIBiG accession BGC0001769) (19), but lacked the *sqnAA*, *sqnA*, and *sqnV-Y* genes (**Supplementary Table 2, Figure 1B**). To confirm that this cluster was indeed responsible for the new antibiotic activity seen for WAC07094, we created insertion mutations in key polyketide synthase genes within the cluster (*sqnH* and *sqnI*; **Figure 1B** and **Supplementary Table 2**) and in parallel, overexpressed the predicted regulatory gene *sqnR*. The mutant strains exhibited a complete loss of antibiotic activity, while overexpressing the SqnR regulator led to enhanced antibiotic activity (**Figure 1C**).

**Figure S3.**
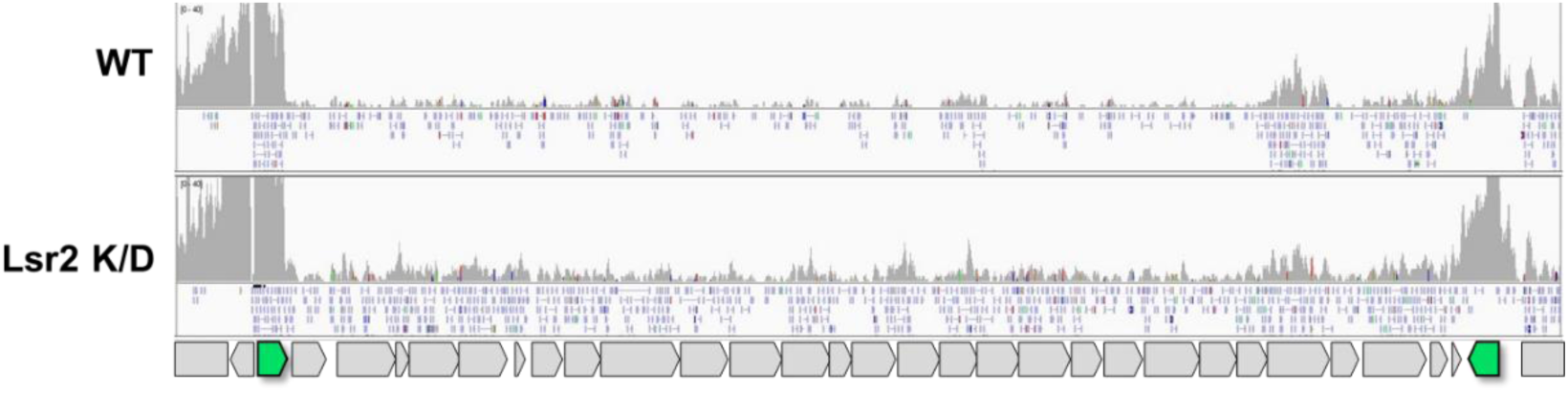
Relative transcript levels for the saquayamycin biosynthetic genes in wild type (WT) WAC07094 and the Lsr2 K/D strain. RNA was extracted from strains grown on Bennett’s agar for 3 days. Transcript levels are represented by RNA sequencing read coverage (grey graphs). Representative sequencing reads are depicted in blue boxes under each graph, for genes oriented in the forward direction. Two regulatory genes (shown as green arrows) are located at either end of the gene cluster.

In parallel, we analyzed the metabolic profiles of crude extracts from wild type and Lsr2 knockdown strains using liquid chromatography and mass spectrometry (LC/MS). We found saquayamycin isomers A and B (with M-H ions at *m/z* 819.29 with a formula of C43H48O16) were readily detectable in the knockdown strain (**Figure 1D**).

### Cell density impacts saquayamycin production and activity

When conducting activity assays for the saquayamycin producer strain, we were surprised to observe that antibiotic activity was only associated with agar plugs taken from the edge of a lawn of WAC07094, and not with plugs taken from the centre. This suggested that saquayamycin production might subject to spatial control, with antibiotic production only being activated at the colony edge, and/or suppression only occurring at the colony centre (**Figure 2A**). To assess whether this was a cell density-dependent phenomenon, we plated a spore dilution series, and evaluated the resulting growth (including both individual colonies and more confluent areas) for antibiotic production using an activity bioassay. We found that activity against *B. subtilis* was inversely proportional to spore density, with increasing colony numbers and greater confluence being associated with reduced antibiotic production (**Figure S4**).

**Figure 2.**
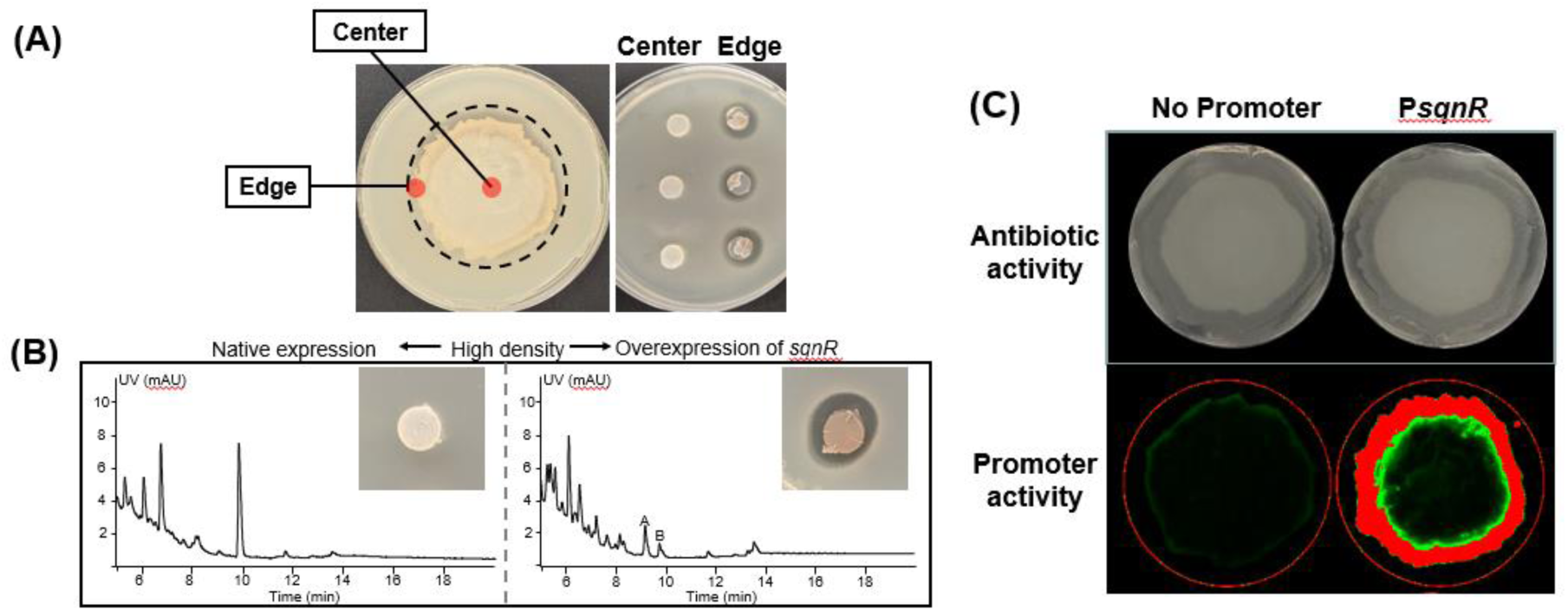
Saquayamycin activity and expression is density-dependent. **(A)** Left: Agar plugs were excised from the center and edge of a confluent lawn of the WAC07094 Lsr2 knockdown strain. Right: Bioassays for antibiotic production, conducted by removing agar plugs (from the colony centre or edge, as indicated on the left) of the Lsr2 knockdown strain, placing them onto *B. subtilis*-containing medium and incubating the plates overnight at 37°C. **(B)** Chromatograms (UV) of crude extracts prepared from Bennett’s agar on which the Lsr2 knockdown strain (left) and same strain carrying the *sqnR* overexpressing construct (right) had been grown as a lawn for 3 days. Peaks of saquayamycin A and B are indicated with the A and B labels, respectively. Inset: anti-*Bacillus* activity exerted by corresponding wild type (left) and *sqnR*-overexpressing (right) strains, where their associated agar plugs were taken from the centre of a confluent lawn (high density). **(C)** Top: Antibiotic bioassay of the Lsr2 knockdown strain carrying either the promoter-less *gfp* (left) or *sqnR* promoter-driven *gfp* construct, against *B. subtilis*. Zones of clearing indicate antibiotic activity. Bottom: promoter activity is seen as green fluorescence produced by the same *Streptomyces* strains, where red signals represent saturated fluorescence intensities (high promoter activity).

**Figure S4.**
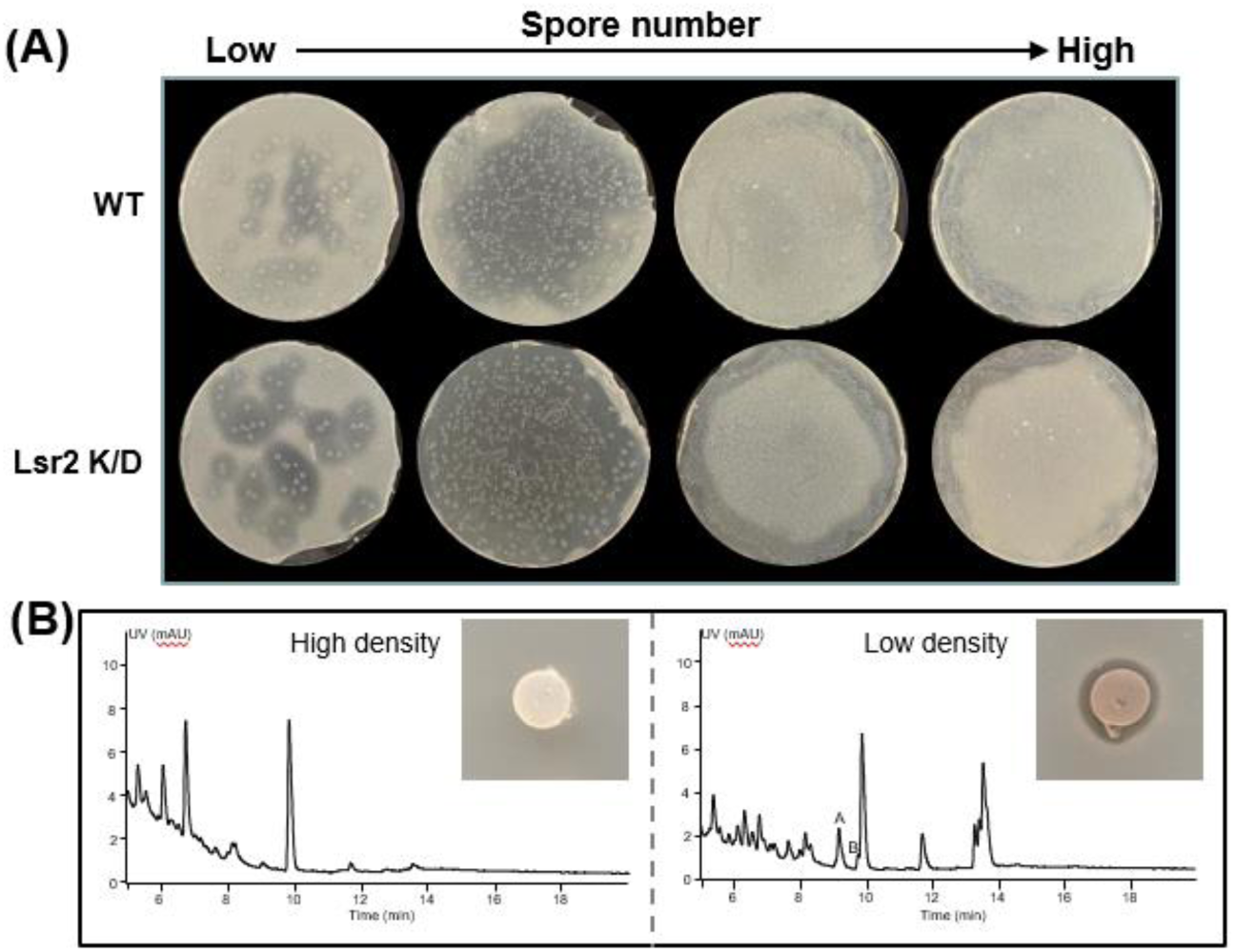
Impact of spore density on antibiotic activity of WAC07094. **(A)** Antibiotic bioassay of a spore dilution series for wild type (WT) and Lsr2 knockdown (K/D) strains against *B. subtilis.* **(B)** Chromatograms (UV) of crude extracts prepared from Bennett’s agar from high-density (left) and low-density (right) Lsr2 knockdown strains grown for 3 days. Peaks corresponding to saquayamycin A and B are indicated with the A and B labels (respectively) within the chromatogram. Inset: anti-*Bacillus* assay using plugs taken from the centre of a confluent lawn (high density) or plugs taken from the centre of a spotted ∼7 mm colony (6 spots/plate to represent a low density condition).

To confirm that the enhanced activity observed during low-density growth was due to saquayamycin, metabolites were extracted from agar medium associated with either high-density or low-density cells. Saquayamycins were detected exclusively in extracts from low-density growth areas (**Figure S4**), and we found this density-dependent effect was alleviated when the pathway-specific regulator *sqnR* was overexpressed (**Figure 2B**), suggesting that the spatial control of antibiotic production may be mediated through *sqnR*. To test this hypothesis, we created a transcriptional reporter fusion using the *sqnR* promoter and a promoterless green fluorescent protein (GFP)-encoding gene. The resulting construct, in parallel with the associated promoter-less control, were introduced into the Lsr2 knockdown strain where they integrated into the chromosome at a heterologous site. We followed the expression of the P*sqnR-gfp* reporter during growth as a lawn and found that GFP expression was only observed at the lawn periphery and was directly correlated with the sites of antibiotic activity (**Figure 2C**).

### High cell-density repression of saquayamycin is alleviated by the addition of magnesium

To begin to understand the basis for the density/spatial specificity of *sqnR* expression, and correspondingly saquayamycin activity, we first considered the effect of nutrients. We tested the impact of various carbon and nitrogen sources, as well as trace metals and salts, on saquayamycin production. We found that magnesium supplementation alleviated the density-dependent suppression of saquayamycin production, with antibiotic production now being observed across the entire colony area (**Figure 3A**). This effect seemed specific to magnesium, as neither zinc nor manganese had an equivalent effect (**Figure S5**). We further tested whether the magnesium-dependent antibiotic stimulatory effect was mediated through *sqnR* using our fluorescent reporter constructs. We found that magnesium supplementation led to *sqnR* expression throughout the colony, again correlating with antibiotic activity (**Figure 3B**). These results suggested that *sqnR* expression, and correspondingly saquayamycin production, could be stimulated by magnesium in a density-independent manner.

**Figure 3.**
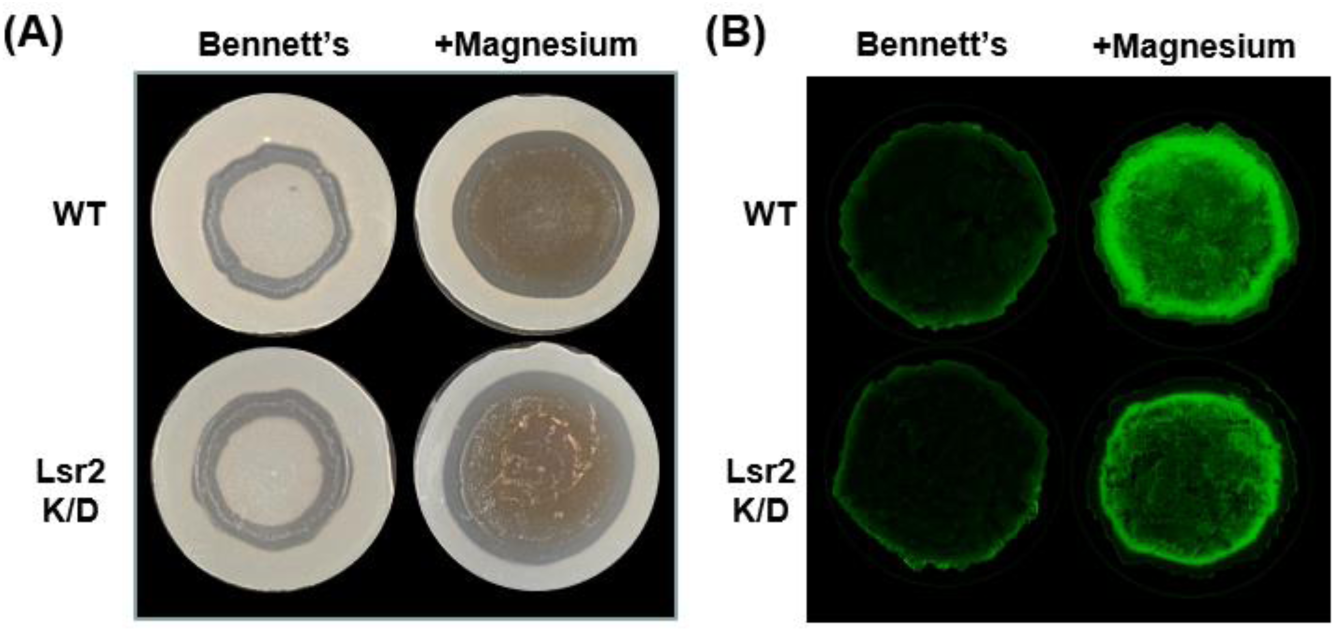
Density-dependent repression of saquayamycin production is alleviated by magnesium supplementation. **(A)** Antibiotic bioassay of wild type (WT) and Lsr2 knockdown (K/D) strains (carrying the *sqnR* transcriptional reporter) grown for 3 days on Bennett’s medium in the absence (left) or presence (right) of magnesium supplementation, where the resulting lawns of the two strains were overlaid with soft agar infused with *B. subtilis.* Zones of clearing indicate antibiotic activity. **(B)** *sqnR* promoter activity of the strains shown in (A), as measured by green fluorescence of different biological replicates.

**Figure S5.**
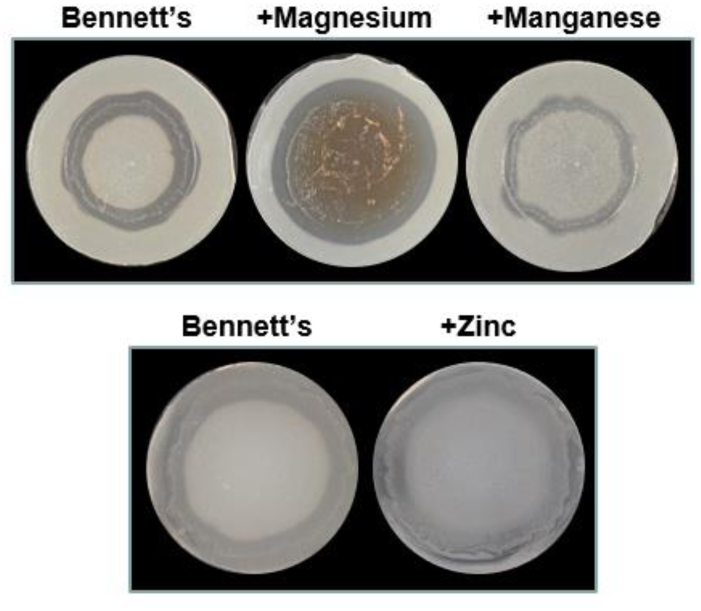
Effect of diverse metal ions on saquayamycin production. Antibiotic bioassay of the Lsr2 knockdown strain grown for 3 days on Bennett’s agar alone, or supplemented with magnesium chloride, manganese chloride, or zinc chloride. Sensitive indicator strain is *B. subtilis*.

### Magnesium and PhoRP independently control saquayamycin biosynthesis

In addition to the effect of nutrients from the environment, we considered the possibility that saquayamycin production might be also governed by quorum sensing. We searched the genome of WAC07094 for genes that may direct the synthesis of extracellular signaling molecules, including butyrolactone and butanolide synthases (commonly used by streptomycetes to synthesize gamma-butyrolactone/butanolide signaling molecules), as well as small peptides like those employed in quorum sensing by other Gram-positive bacteria. Unexpectedly, no strong candidates for genes encoding signaling peptides or butyrolactone/butanolide synthases were found in the genome.

When searching for small peptide-encoding genes, however, we did identify one termed *mtpA* that was predicted to encode a small metal-binding protein (20), and wondered whether there may be a connection between the resulting MtpA protein, magnesium, and saquayamycin production. To localize *mtpA* expression, we constructed a transcriptional fusion between the *mtpA* promoter and the promoterless *gfp* reporter gene, and introduced the resulting construct into the wild type strain, where it integrated into the chromosome at a heterologous site. We found that like *sqnR, mtpA* expression was enhanced at the colony periphery and was increased throughout the colony following magnesium (but not manganese or zinc) supplementation (**Figure 4A**).

**Figure 4.**
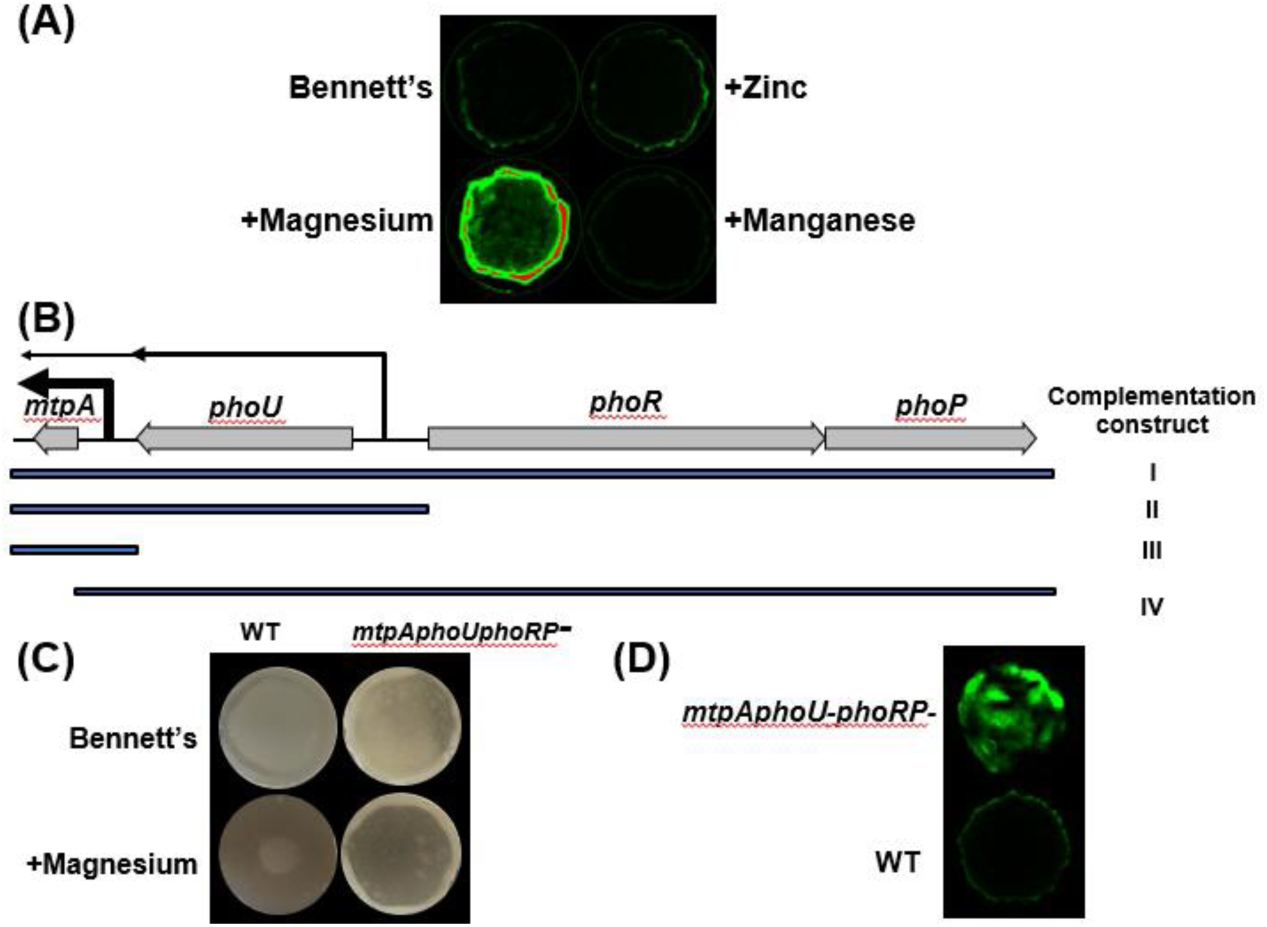
The density-dependent expression of *mtpA,* and the saquayamycin cluster is affected by magnesium and PhoRP. **(A)** Promoter activity of *mtpA* in wild type WAC07094 after 3 days growth on Bennett’s agar with or without magnesium chloride, manganese chloride or zinc chloride supplementation. **(B)** Genetic organization of the *mtpA*, *phoU*, *phoR*, and *phoP* locus. Transcription start sites are indicated by vertical lines, with the line widths (and associated horizontal arrows) approximating relative transcript abundance. Beneath the genes are four lines, indicating the sequence included in each of four complementation constructs (labelled I to IV on the right). **(C)** Antibiotic activity as detected using *B. subtilis* infused agar, overlaying lawns of wild type (WT) or mutant strains grown for 3 days on Bennett’s agar, with or without additional magnesium supplementation. **(D)** Promoter activity of *sqnR* after 3 days of growth, as indicated by the relative fluorescence within the lawn of the wild type and *phoRP* locus mutant.

We noted that *mtpA* was located immediately downstream of *phoU* (**Figure 4B**). Our RNA-seq data suggested that *mtpA* was expressed primarily from its own promoter, but that read-through transcription from *phoU* also contributed to its expression (**Figure 4B**). These two genes in turn, were divergently oriented from the *phoRP* two-component system-encoding genes (**Figure 4B**). To determine whether there were connections between any of these genes/proteins and saquayamycin production, the four gene (*mtpAphoU-phoRP*) locus was deleted in the wild type background, and both *sqnR* expression and antibiotic activity were monitored. Interestingly, the mutant phenotype was reminiscent of colonies growing with magnesium supplementation, although not quite as robust: antibacterial activity and *sqnR* expression were now detected throughout the colony, not just at the colony edges (**Figure 4C** and **4D**, respectively). This suggested that the density-dependent saquayamycin production phenotype could be facilitated at least in part by one or more of the genes in the *mtpAphoU-phoRP* locus. We also observed that the loss of *mtpA* (as part of this four gene locus) did not impact the magnesium-mediated stimulation of saquayamycin production **(Figure 4C**), suggesting that this effect did not require MtpA activity.

To determine which of the gene products within this locus was responsible for the density-dependent saquayamycin biosynthesis phenotype, we constructed a suite of complementation constructs (**Figure 4B**) which were introduced into the *mtpAphoU-phoRP* mutant strain. We first confirmed that complementation with the four-gene locus restored the wild type phenotype, with antibiotic production only being observed at the colony periphery (**Table 1**). Complementation with *phoRP* together with *phoU* (lacking *mtpA*), had an identical effect, with saquayamycin activity being restricted to the colony periphery (**Table 1**). This suggested that MtpA was not responsible for the spatial localization of saquayamycin production. In contrast, when *mtpA* and *phoU* were re-introduced (without *phoRP*), colony-wide antibiotic production was retained. This indicated that the spatial nature of saquayamycin production was likely due to PhoRP-mediated repression of *sqnR* expression in high cell density areas.

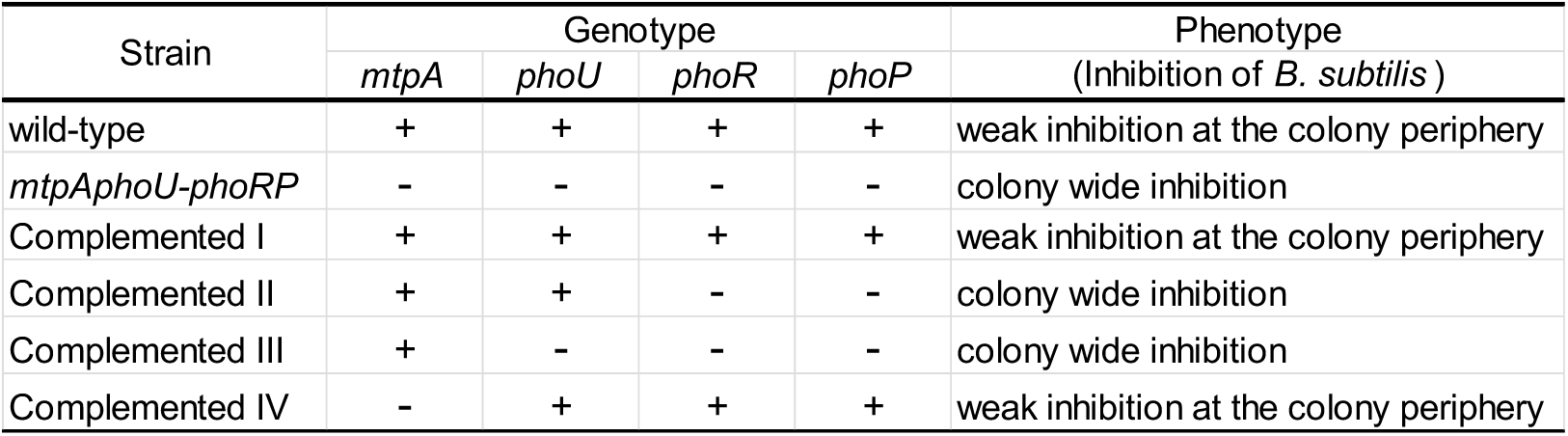
Antibiotic activity phenotype of the mtpAphoU-phoRP complementation strains.

### Saquayamycin production is impacted by multiple regulators

The Pho system controls phosphate transport, and the PhoRP system is activated in response to low phosphate levels (11). This results in phosphorylation of the PhoP response regulator, which enables its binding to the promoters of target regulon members at well-defined Pho-box sites (21–22), altering the expression of the associated genes. Given the negative effect of PhoRP on *sqnR* expression – and saquayamycin production – in high density colony areas (**Figures 4C** and **4D**), we examined the *sqnR* promoter for possible PhoP binding sites. We identified a candidate PhoP-box located 28 nt upstream of the predicted transcription start site of *sqnR* within the likely promoter region (**Figure 5A**). An equivalent box (direct repeat units GG/TTCAYYYRC/GG [21]) was also found upstream of *phoU.* Beyond PhoP, the binding sites of many other global regulators of antibiotic production have been defined, and a number of these share core recognition sequences with PhoP, including GlnR and the two component regulators MtrA and AfsQ1 (**Figure 5A**) (9), and thus may also impact *sqnR* expression and saquayamycin production. We overexpressed each regulator-encoding gene (for *afsQ1,* we overexpressed a mutant version expressing a phosphomimic variant [23]) to determine whether any of these impacted saquayamycin production in either wild type or Lsr2-knockdown backgrounds. As expected, enhanced antibacterial activities were observed for all Lsr2 knockdown-carrying constructs (**Figure 5B**). We found that overexpressing *mtrA* (together with its associated kinase-encoding *mtrB*) reduced saquayamycin production, while overexpressing *glnR* and *afsQ1\** [from constitutive (*ermE\**) and inducible (*tipA*) promoters, respectively] enhanced saquayamycin production in both wild type and Lsr2 knockdown backgrounds (**Figures 5B, 5C)**. In the case of *afsQ1\**, its effects were most pronounced in the wild type background, with Lsr2 knockdown having no additive effect in an *afsQ1*\* overexpressing strain (**Figure 5B**). This suggested that AfsQ1 may function to counteract the repressive function of Lsr2. In contrast, overexpressing *glnR* in the wild type background was at least as effective as Lsr2 knockdown in enhancing saquayamycin production, while in the Lsr2 knockdown background, the effects were additive and mirrored the effect of *phoRP* deletion. This collectively suggested that PhoP and Lsr2 were the predominant repressors of saquayamycin production, while GlnR and AfsQ1 contributed to saquayamycin activation.

**Figure 5.**
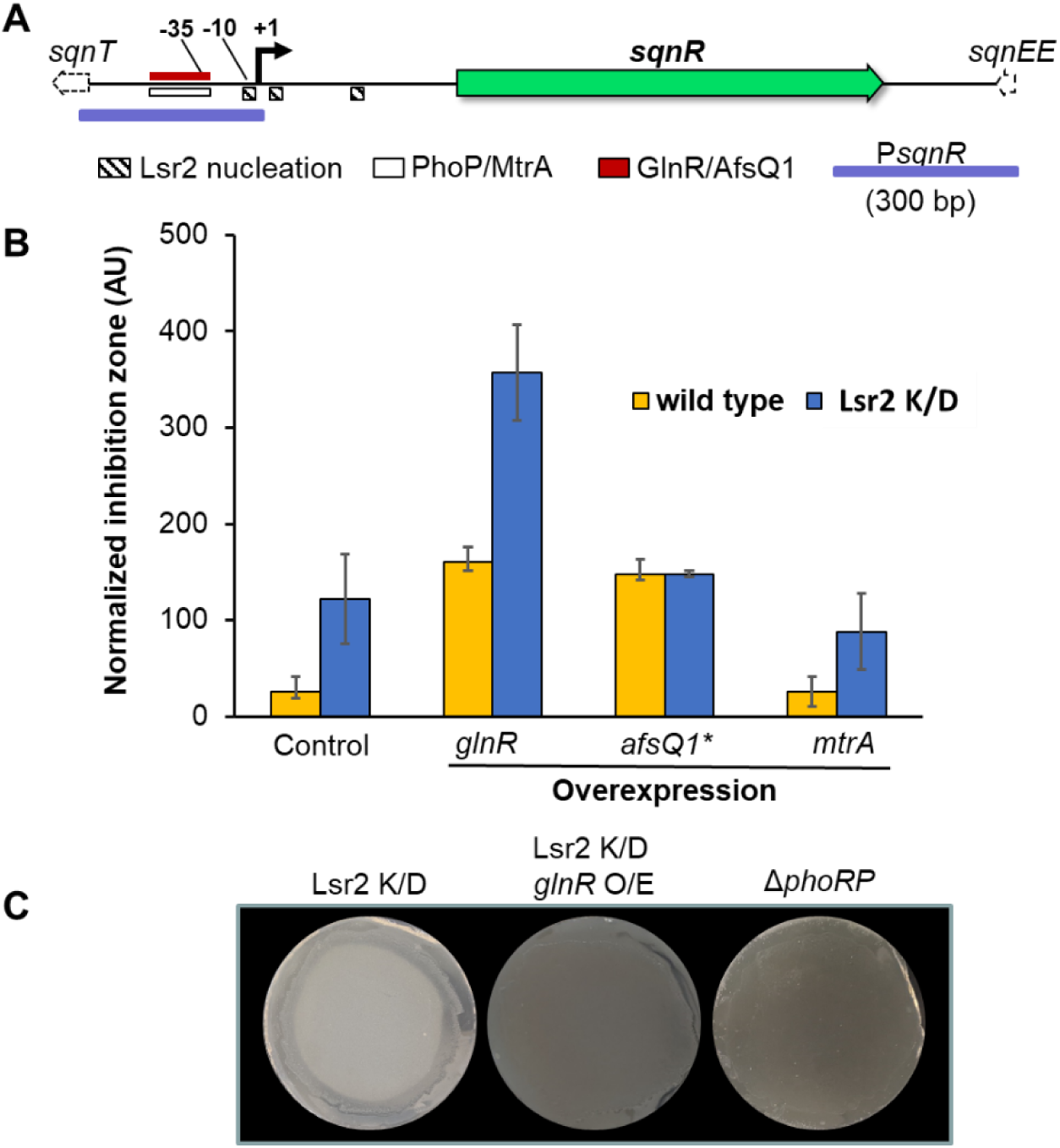
GlnR positively controls saquayamycin production. **(A)** Schematic of the *sqnR* promoter region, including the transcription start site (+1), the upstream promoter (-10 and -35), and the predicted binding sites for Lsr2, PhoP, MtrA, GlnR and AfsQ1. The promoter region used for the reporter constructs is indicated in purple. **(B)** Antibiotic bioassay using WAC07094 wild type (WT) or Lsr2 knockdown (K/D) strains carrying either the empty plasmid pIJ10257 (Control), or *glnR* or *mtpA* under the highly active *ermE\** promoter, or *afsQ1\** under the inducible *tipA* promoter. *B. subtilis* was used as the sensitive indicator strain, and growth inhibition zones were quantified and normalized to the corresponding *Streptomyces* growth area. Error bars indicate standard deviation (*n*=4). **(C)** The effect of *phoRP* deletion relative to Lsr2 knockdown, and Lsr2 knockdown together with *glnR* overexpression (O/E), were assessed using *B. subtilis*-infused agar overlays. The darker coloration associated with the two right-most strains indicate full growth inhibition, in contrast to inhibition only at the periphery seen for the left-most plate.

### Saquayamycin biosynthesis is activated in low-density inocula liquid cultures

Given the intriguing density-dependent production of saquayamycin during growth on solid agar medium, we wondered whether an equivalent density-dependence might also be observed for liquid-grown cultures. We tested the antibiotic production activity of both wild type and Lsr2 knockdown strains using high and low concentration spores as inocula in liquid Bennett’s medium. We found that both strains produced saquayamycin exclusively in cultures inoculated with low numbers of spores (**Figure 6A**). We tested whether magnesium supplementation or *phoRP* deletion could enhance saquayamycin production when starting with a higher spore inoculum. We found that unlike on solid medium, magnesium failed to stimulate saquayamycin production during ‘higher density’ (higher spore inoculum) growth. In contrast, *phoRP* mutants produced high levels of saquayamycin under the same growth conditions (**Figure S6**).

**Figure 6.**
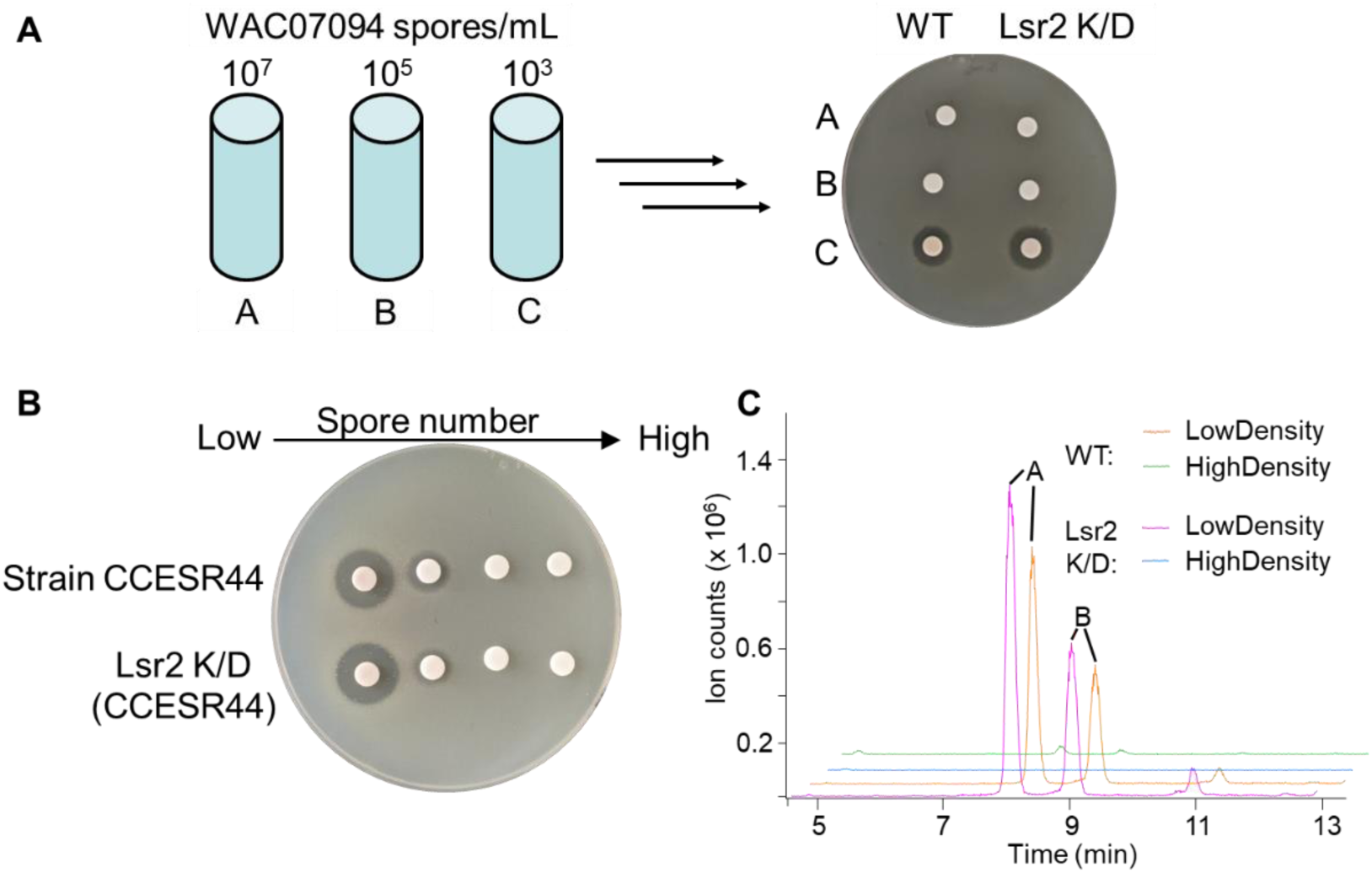
The density-dependence of saquayamycin production is conserved across diverse streptomycetes. **(A)** Antibiotic bioassays were conducted using *B. subtilis* as the sensitive indicator organism, together with extracts of wild type (WT) and Lsr2 knockdown (K/D) WAC07094 strains cultures inoculated from low (10^3^/mL), mid (10^5^/mL) and high (10^7^/mL) density spore preparations and grown for 4 days in Bennett’s liquid medium. **(B)** Equivalent experiments to that in (A) were conducted for *Streptomyces* isolate CCESR44 wild type and Lsr2 knockdown strains, only including an additional low-density inoculum of 100-200 spores/mL. **(C)** Extracted ion chromatograms of [M-H]^-^ (at *m/z* 819.3) from metabolite profiles for low and high cell-density inoculated cultures from (B).

**Figure S6.**
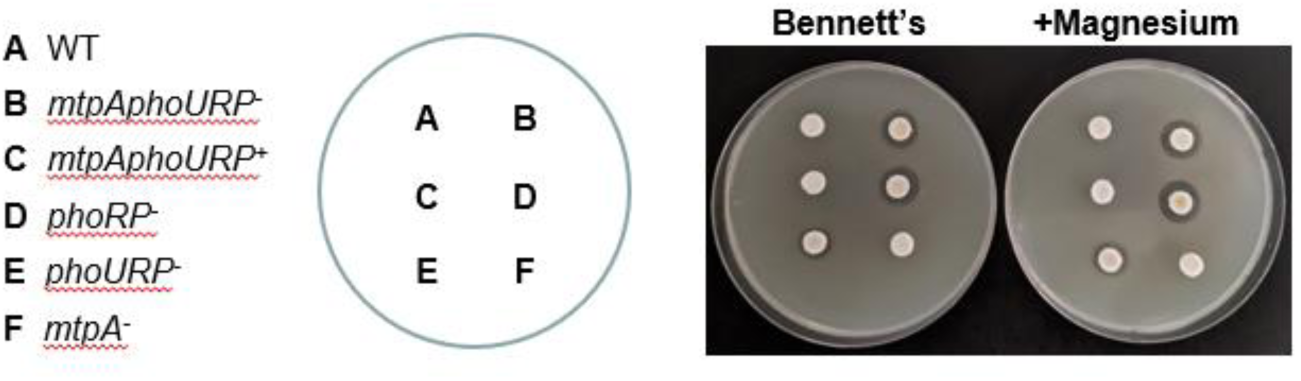
Loss of PhoRP can overcome the density-dependent production of saquayamycin during liquid culture growth. Antibiotic bioassays were conducted using extracts from high-density liquid cultures of WAC07094 (inoculated with 10^7^ spores/mL) grown for 4 days in Bennett’s liquid medium. Ten microlitres of extract from each of the six wild type and mutant cultures were applied to filter discs overlaid on medium mixed with *B. subtilis*.

### Control of saquayamycin production is conserved between different producer strains

To investigate whether the low cell density production phenotype was shared more broadly amongst saquayamycin producers, we searched for strains with this biosynthetic cluster using the National Center for Biotechnology Information (NCBI) and Joint Genome Institute (JGI) databases (24–25) and found a candidate in *Streptomyces* sp. 3212.3 (Accession no. QTTM01000001.1), which has subsequently been termed isolate CCESR44 (L. Kinkel, personal communication). We compared its annotated saquayamycin biosynthetic cluster to that of WAC07094 and found only minor differences at the cluster boundaries (**Figure S7A**). Phylogenetic comparisons between these strains and several reference genomes (de novo mode in https://automlst.ziemertlab.com/index) (26) were conducted using ∼50 select single copy conserved genes (**Figure S7B**, supplemental text file). These analyses suggested that strains WAC07094 and CCESR44 were unique, but shared 99% identity shared between 16S rRNA sequences (comparing all 6 copies of the gene); they grouped closely with each other and with *Streptomyces* sp. NRRL F-5122 and *Streptomyces nodosus* ATCC 14899. Strains WAC07094 and CCESR44 were also observed to have different colony morphologies (**Figure S7C**).

**Figure S7.**
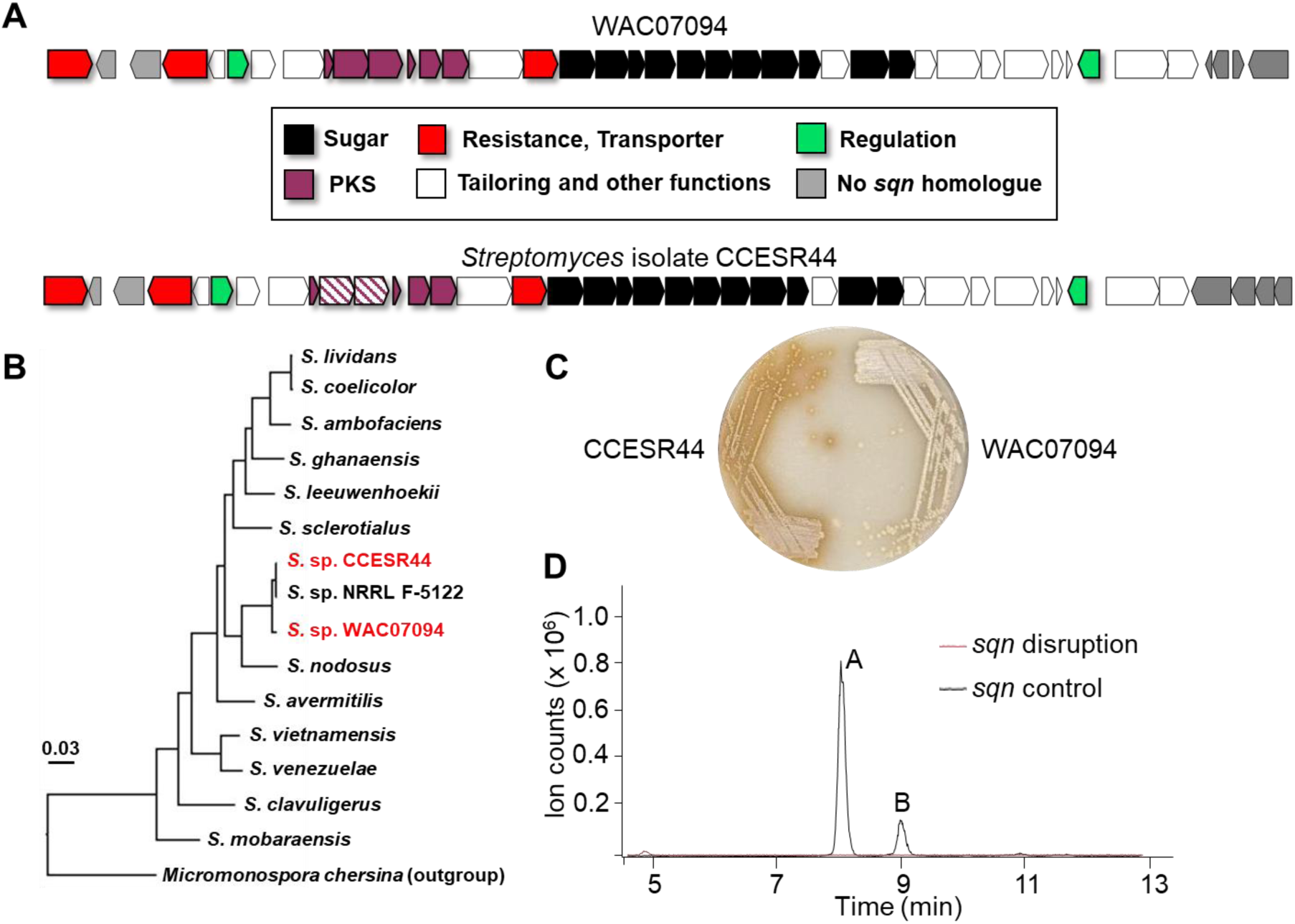
Comparison of saquayamycin producer strains WAC07094 and *Streptomyces* isolate CCESR44. **(A)** Saquayamycin biosynthetic clusters in WAC07094 and the alternative producer *Streptomyces* isolate CCESR44. Minor differences of gene organization are coloured in grey. Those biosynthetic genes targeted for disruption (by homologous recombination) are indicated in purple hatching. **(B)** Phylogenetic tree of strains WAC07094 and CCESR44 relative to other streptomycetes, using *Micromonospora chersina* as an outgroup. Bootstrap values of each branch: greater than or equal to 50% (based on 1000 resampled trials). The scale bar (labelled 0.03) represents substitutions per nucleotide position. The phylogenetic analysis was conducted using https://automlst.ziemertlab.com/index (26) based on 50 select single copy conserved genes. **(C)** Phenotypic comparison of WAC07094 and CCESR44 following growth on mannitol-soy flour agar. **(D)** Extracted ion chromatograms of [M-H]^-^ (at *m/z* 819.3) from the extracts of the 3212.3 *sqnHI* mutant and WAC07094 (as a positive control) grown from low-density spore inoculum in Bennett’s liquid medium for 4 days.

We tested the effect of cell density on saquayamycin production by CCESR44. Interestingly, this strain was also observed to exclusively produce saquayamycin when grown using a low density inoculum in liquid culture (**Figure 6B**). Introducing our Lsr2 knockdown construct into this strain further increased antibiotic activity (**Figure 6B** and **6C**). We confirmed that this activity was due to saquayamycin using chemical analyses, alongside genetic tests, where creating insertion mutations in the polyketide synthase genes *sqnH* and *sqnI* abolished antibiotic activity (**Figure S7D; Figure 6C**). We further tested the effects of magnesium supplementation during growth on solid medium, and as for WAC07094, we observed antibiotic production throughout the entire colony, irrespective of cell density (**Figure S8**). These observations collectively suggest that the spatial, genetic and nutritional controls governing saquayamycin production are conserved across disparate species.

**Figure S8.**
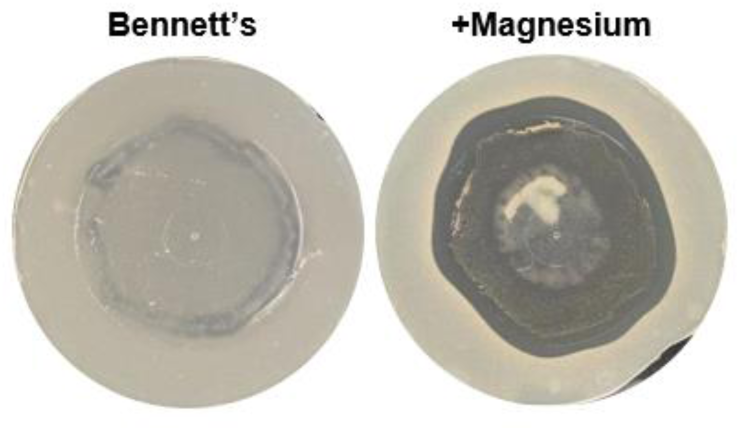
Magnesium supplementation can overcome the density-dependent saquayamycin production of strain 3212.3. Antibiotic bioassays using *B. subtilis* as the indicator strain inoculated into agar that was overlaid atop *Streptomyces* sp. CCESR44 grown on Bennett’s agar for 3 days with and without magnesium supplementation.

## DISCUSSION

The conventional view of specialized metabolism in the streptomycetes is that it occurs in mature, high-density cultures. Our work here suggests that there are conserved exceptions to this rule, where antibiotic production can instead be confined to low cell density areas. In bacteria, cell density can impact a wide range of processes (27–29). One of the best-studied density-dependent behaviours involves quorum sensing, which in *Streptomyces* species is mediated predominantly by butyrolactone or butanolide signalling molecules (30). While there were no obvious quorum signalling systems encoded in the genome of WAC07094, our work suggests that this species effectively exerts density-dependent control of the saquayamycin biosynthetic genes via the cluster-situated regulator *sqnR*.

We found that there were multiple factors that influenced the density-dependent production of saquayamycin (**Figure 7**). Magnesium was a particularly potent activator of *sqnR* expression during solid culture growth. We were not able to distinguish whether this was the result of specific induction of *sqnR* transcription, or if it reflected a broader impact on cellular physiology; however, it is worth noting that the saquayamycin-enhancing effect of magnesium was conserved in both *Streptomyces* species WAC07094 and CCESR44. In other bacteria, magnesium has been implicated in everything from DNA replication (31) and gene regulation (32–34), through to protein folding and stability (35). It has also been associated with alternative oligomeric assemblages for the nucleoid-associated protein H-NS (36), which is functionally equivalent to the Lsr2 protein in the actinobacteria. Whether magnesium impacts Lsr2 behaviour is currently unknown; it is tantalizing to speculate that it may help to alleviate the Lsr2-mediated repression observed for the saquayamycin biosynthetic cluster.

**Figure 7.**
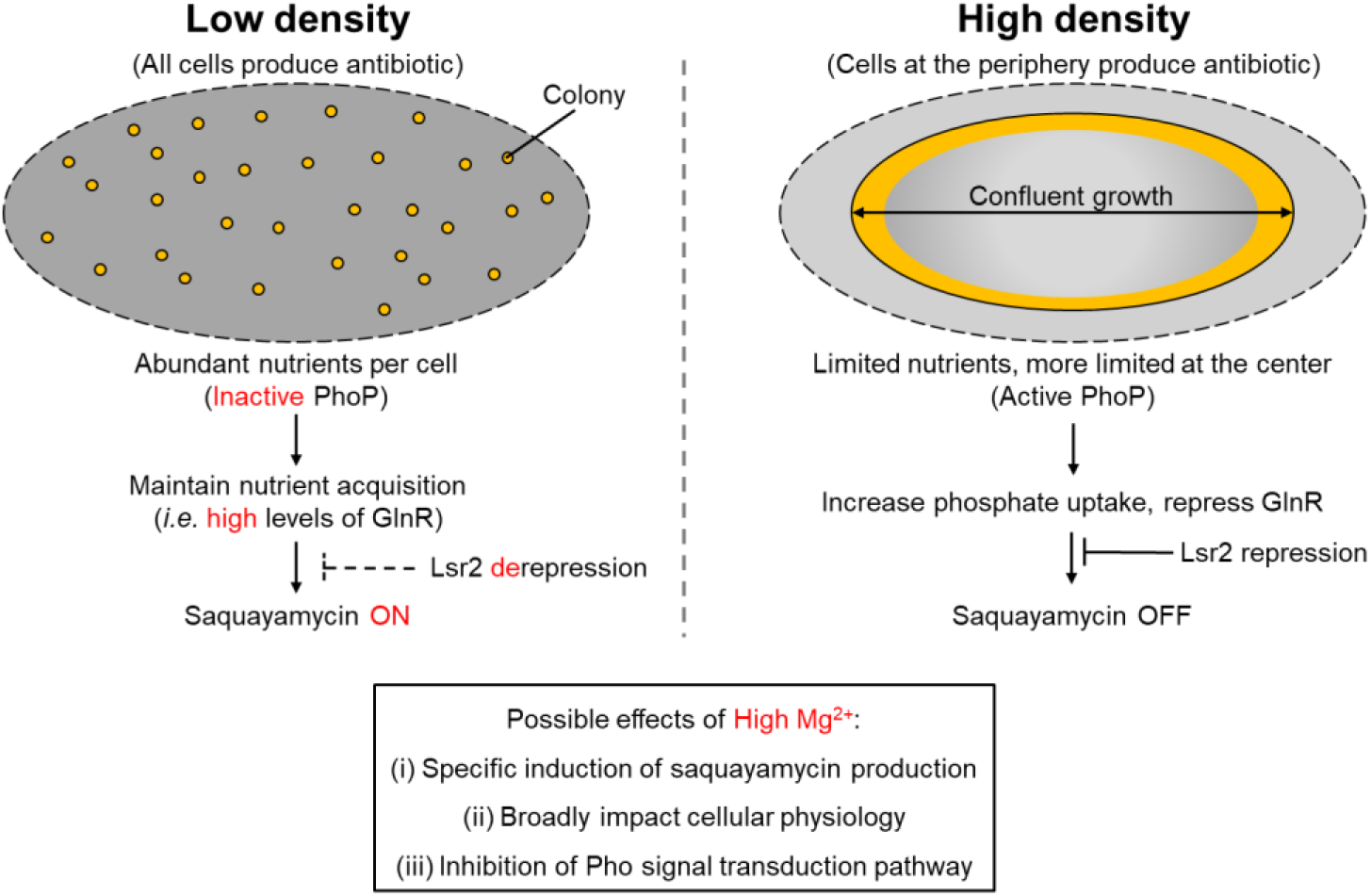
Model of the regulatory and nutritional inputs impacting saquayamycin production.

The effects of magnesium on gene regulation have been best studied in the enterobacteria (*Salmonella, Yersinia, Escherichia*), where magnesium levels modulate the activity of the PhoP/PhoQ two-component regulatory system (37). The streptomycetes do not have an equivalent system; instead, they and their Gram-positive relatives possess a PhoP/PhoR system, which is analogous to PhoB/PhoR in the enteric bacteria. In *Salmonella,* increased magnesium levels also negatively affect RNA polymerase occupancy and mRNA transcript levels of PhoB regulon members, including genes for the phosphate transport system (*pstSCAB*), and *phoBR* itself (38). While the effect of magnesium on PhoP/PhoR activity in the streptomycetes is unknown, a similar effect could lead to reduced PhoP levels. Notably, deletion of *phoRP* led to constitutive *sqnR* production and enhanced saquayamycin production, similar to the phenotype observed following magnesium supplementation. This suggested that PhoP functioned to repress *sqnR* expression in areas of confluence/high cell density, and was further supported by the presence of a Pho box upstream of the *sqnR* transcription start site. We cannot, however, exclude the possibility that PhoP controls the synthesis of a novel quorum (or anti-quorum) sensing compound that impacts *sqnR* expression. There is precedent for connections between phosphate regulation and quorum sensing in *Pseudomonas aeruginosa*, where production of the IQS signalling molecule is governed by PhoB, the PhoP equivalent (39).

In the absence of PhoP repression, we propose that transcription activator(s) can access the *sqnR* promoter and recruit RNA polymerase to help counter the downstream Lsr2 silencing of the saquayamycin gene cluster (40). A likely candidate for this role is the nitrogen-responsive GlnR regulator. There is a predicted GlnR binding site upstream of the *sqnR* promoter, and overexpression of GlnR led to increased saquayamycin production. There is further an intriguing reciprocal relationship shared by GlnR and PhoP in many streptomycetes (9) that mirrors the situation with saquayamycin production. PhoP can directly repress GlnR, and thus as PhoP levels drop, GlnR levels would rise. In WAC07094, this could allow GlnR to both activate *sqnR* transcription, and promote the RNA polymerase-mediated removal of Lsr2 from the chromosome, alleviating its repressive effects. AfsQ1, which shares the same DNA target sequence with GlnR, can also promote saquayamycin production, and this appears to be largely through alleviation of Lsr2 repression.

The ecological roles of most natural products are not well understood. Saquayamycins are pigmented, glycosylated natural products, and they have growth-inhibitory activity against both Gram-positive bacteria and eukaryotes. Indeed, the activity of the saquayamycins and other angucycline family members have been best studied for their antitumor and enzymatic inhibitory properties (41). Paradoxically, angucyclines have been reported to be produced by endophytes – microbes that live inside plants and sponges (42–43). We speculate that the producers of these compounds, and in the case of endophytes, also their hosts, must benefit from the synthesis of these toxic compounds, possibly as a result of their antifungal properties. In the case of free-living bacteria like WAC07094 and CCESR44, the production of saquayamycin at the colony edge may function to stave off competing microbes and secure sufficient nutrients to promote continued growth. It is notable that saquayamycins are synthesized in many forms (here we identified saquayamycins A and B). Whether different forms have distinct functions or properties remains to be determined.

Microbes are an extraordinary source of biologically active molecules. Understanding the interplay between gene regulation, physiology and metabolism is important to maximize the biosynthetic capacity of these organisms, and to exploit the functional landscape of new molecules. Beyond demonstrating the power of our Lsr2 knockdown approach in stimulating saquayamycin production, and uncovering the inter-related impacts of magnesium, PhoP, GlnR, and AfsQ1, our work here has also revealed the unexpected importance of low cell density on specialized metabolism. This observation is not without precedent: in 2011, a blue pigmented molecule was found to be produced exclusively at lower initial cell densities by the plant pathogenic bacterium *Pantoea agglom*erans (44). As this earlier report suggests, this phenomenon is unlikely to be confined to specific angucyclines, and our findings here suggest there may be benefits to adding low cell density cultures to antibiotic screening platforms.

## MATERIALS AND METHODS

### Bacterial strains, plasmids, and culture conditions

Bacterial strains, plasmids, and oligonucleotides used in this study are listed in Supplementary Tables 3-4. *E. coli* DH5α was used for general subcloning and plasmid preparation (45). *E. coli* strains were grown at 37°C in lysogeny broth (LB) and Difco™ nutrient agar. The non-methylating *E. coli* strain ET12567/pUZ8002 was used as a donor host for intergeneric conjugation (46). *Streptomyces* strains were grown at 30°C in liquid Bennett’s medium or on Bennett’s agar (per L, 10 g glucose, 1 g yeast extract, 1 g beef extract, 2 g N-Z amine, without or with 15 g agar) (47) or cultured on International *Streptomyces* Project 4 agar (ISP4) (48) unless otherwise stated. Antibiotic selection was carried out using antibiotics at the following concentrations: apramycin (50 μg/mL), chloramphenicol (25 μg/mL), kanamycin (50 μg/mL), nalidixic acid (25 μg/mL) and hygromycin (50 μg/mL). The indicator strains used in the antimicrobial assays included *Bacillus subtilis*, methicillin-resistant *Staphylococcus aureus*, and vancomycin-resistant *Enterococcus faecium*. These strains were grown at 37°C in Tryptic Soy Broth or on Difco™ nutrient agar.

### Whole genome sequencing

Genomic DNA was prepared using a salting out protocol (47), after which RNA was removed using an RNase A treatment. DNA quality and quantity were assessed using agarose gel electrophoresis and a Nanodrop spectrophotometer (ThermoFisher). A draft genome sequence for *Streptomyces* sp. WAC07094 was obtained using both Illumina and PacBio sequencing. For Illumina sequencing, library preparation, sequencing, and read processing were conducted as described previously (49). For PacBio sequencing, the DNA library was prepared by the Farncombe Metagenomics Facility at McMaster University using the SMRTbell Template Prep Kit 1.0 following PacBio’s Microbial Multiplexing protocol. Multiple libraries were pooled and size-selected using the BluePippin BLF75 kit with a high pass collection for fragments >7 kb. Sequencing was performed in the Farncombe Metagenomic sequencing facility using the Sequel Sequencing Kit 3.0 reagents for a 20 hour-run on a SMRT Cell 1M. PacBio reads were assembled using the Flye assembler (50) in a computer cluster before being polished first with the built-in Flye system, followed by the gccp polisher (https://github.com/PacificBiosciences/gcpp). A ‘reference’ sequence was aligned with PacBio raw reads using pbmm2 aligner (https://github.com/PacificBiosciences/pbmm2). The polished assembly was ultimately aligned with filtered and trimmed Illumina reads using minimap2 aligner (51). The overlapping sequences were used to create a consensus sequence between the Illumina and PacBio sequencing reads using Racon (52), to give a final assembly consisting of four contigs (8,469,185; 263,838; 536,999; 329,400 bp). The contig sequences have been deposited in DDBJ/ENA/GenBank under the accession JAVMJZ000000000. The version described in this paper is version JAVMJZ010000000. Biosynthetic gene clusters were predicted using antiSMASH 5.0 (18).

### Genetic manipulation of WAC07094

To modulate the activity of Lsr2 in WAC07094, a previously generated Lsr2 knockdown construct (pMC109) (12) was used. This plasmid was introduced into WAC07094 through intergeneric conjugation. About 10^8^ viable spores were used for conjugation on ISP4 supplemented with 20 mM MgCl2 after mixing with cells from 3 mL of *E. coli* ET12567/pUZ8002 overnight cultures. Representative apramycin-resistant exconjugants were selected for bioassays. As a control, the pIJ12551 (empty plasmid parent of the Lsr2 knockdown construct) was also introduced into WAC07094.

To disrupt the saquayamycin biosynthetic genes, a 2.3 kb-homologous DNA fragment was cloned into a unique *Spe*I site of a non-replicative plasmid (pIJ10700) using T4 DNA ligase (Roche). The fragment was amplified using Q5^®^ DNA polymerase (NEB) and primers DRsqnH-upSpeI/DRsqnI-downSpeI with a 61-68°C gradient annealing temperature (Bio-Rad S1000 thermocycler). The cloned fragment was confirmed by restriction analysis and sequencing using the universal M13-Reverse primer. Gel extraction and plasmid preparation kits were purchased from NEB and Invitrogen, respectively. The resulting disruption construct (pMC341) was introduced into the WAC07094 Lsr2 knockdown strain through conjugation and selection for hygromycin resistant exconjugants.

To overexpress the transcriptional activator-encoding *sqnR*, the gene was cloned into the *Nde*I and *Xho*I sites, downstream of the constitutive *ermE\** promoter in the integrating plasmid pIJ10257. *sqnR* was amplified using primers EXsqnR-upNdeI/EXsqnR-downXhoI with a 62-67°C gradient annealing temperature. The resulting plasmid clones (pMC342) were sequenced using primer P*ermE* for verification, before being mobilized into the WAC07094 Lsr2 knockdown strain. As above, the empty pIJ10257 was also introduced into the knockdown strain as a control.

### Antibiotic bioassays

To test their bioactivities, *Streptomyces* sp. WAC07094 strains or strain 3212.3 were inoculated onto Bennett’s agar plates in various ways, including: (i) applying 5 µl of a spore suspension to yield a dense ‘spot’ (*e.g.* **Figure 1**), (ii) spreading spore suspensions (100 µL each) over an agar plate to obtain a lawn (*e.g.* **Figure 2**), or (iii) spreading to obtain single colonies (*e.g.* **Figure S4A**). In each instance, the resulting cells/colonies were grown for 3 to 5 days. The day before initiating the bioassay, indicator strains were inoculated and grown overnight, before being inoculated (0.1% v/v) into melted Difco™ nutrient agar. The resulting cell suspension was then poured into a petri plate. Subsequent antibiotic tests were conducted using either ‘agar plug’ or ‘sandwich’ assays. In the case of the former, agar plugs were removed from the lawn of *Streptomyces* growth and transferred onto the indicator plates. In the latter situation, the indicator-containing agar was removed from its petri dish once solidified and was overlaid atop the *Streptomyces* growth plate. For both plug and sandwich assays, the plates were incubated overnight at 37°C. Antibiotic effects on the indicator strain were observed as inhibition zones around the plugs or in/around the *Streptomyces* growth areas, and these were captured by camera (Pixel 3a). The inhibition zones were measured using ImageJ 1.49v at 8-bit resolution (53), and were normalized against growth area, which was also measured using the same software.

Antibiotic activity of crude extracts (the preparation for which is described below) was determined using 6 mm disc-diffusion assays on indicator plates. The indicator plates were prepared as described above, after which the 6 mm filter discs were place on the plate, and 5-10 µL of crude extract were applied to the discs. Growth inhibition of the indicator strains was observed as zones of clearing around the disks after overnight incubation.

### Genome-wide transcriptome analysis using RNA-Seq

With a genome assembly in hand, we set out to assess the expression of biosynthetic clusters under conditions where bioactivity was observed for the Lsr2 knockdown strain. *Streptomyces* sp. WAC07094 was inoculated by spotting 5 µL of spore suspensions in sterile water (5,000 to 50,000 viable spores) on the Bennett’s agar plates. After 3 days of growth, RNA was isolated from biomass scraped from the agar surface (∼100 mg) using the Macherey-Nagel NucleoSpin^®^ RNA isolation kit with minor modifications. Cells were lysed using beads and chloroform, prior to RNA purification using the spin column. To remove contaminating DNA, RNA samples were treated with TURBO™ DNAse (Ambion), and the resulting RNA preparations were then assessed for abundance and purity using a Nanodrop spectrophotometer, with RNA integrity being further assessed using agarose gel electrophoresis and an Agilent 2100 Bioanalyzer. To ensure there was no residual DNA in the RNA preparations, a housekeeping gene (16S rRNA gene) was targeted for amplification (35 cycles using primers listed in Table S4).

cDNA library construction and sequencing were performed by the Farncombe Metagenomics Facility at McMaster University using an Illumina MiSeq sequencer with a MiSeq v3 reagent kit (2 ×75 bp paired end configuration). Prior to library construction, ribosomal RNA was depleted using RiboZero (Illumina). Libraries were prepared using NEBNext Ultra II Directional RNA library prep kit (without size selection) and barcoded using NEBNext^®^ Multiplex Oligos for Illumina^®^ (96 Unique Dual Index Primer Pairs).

The raw sequencing outputs were filtered and trimmed for adapter sequences using skewer (54). The filtered and trimmed reads were checked for quality control using FastQC (https://www.bioinformatics.babraham.ac.uk/projects/fastqc/) and aligned to their respective genome sequence using Bowtie2 (55). The aligned sequences were visualized using Integrated Genomics Viewer (56).

### Analysis of metabolite extracts

Metabolites were extracted from cultures of *Streptomyces* sp. WAC07094 strains grown on Bennett’s agar or liquid medium. For agar-grown cultures (3 days of growth), agar (including cell biomass) was chopped into small pieces and soaked overnight in ethyl acetate (agar/biomass from two 10 cm plates in 50 mL). The organic solvent was evaporated under reduced pressure using a rotavap (Heidolph) and the resulting pellet was resuspended in 500 µL of methanol:acetone (1:1). The suspension was dried under reduced pressure and resuspended in 500 µL of a 1:1 mixture of methanol and water. For 4-day old submerged cultures, 5 mL of cell-free broth was extracted using ethyl acetate at a 1:1 ratio. The organic phase was then removed and evaporated under reduced pressure using a speedvac (Thermo Scientific). The resulting pellets were dissolved in 50 µL DMSO. After the removal of particulates by centrifugation, the extracts were subjected to LC-MS analyses under UV and ion detection.

Two approaches were used for saquayamycin detection and verification. The first involved analytical HPLC using an Agilent 1290 Infinity LC System with a Zorbax Eclipse XDB C18 column (100 mm × 2.1 mm × 3.5 µm, respectively), coupled to an LTQ Orbitrap XL MS system (Thermo Scientific). Metabolites were separated at a flow rate of 0.4 mL/min using a 2.5 min solvent gradient, from 95% solvent A (water with 0.1% formic acid) and 5% solvent B (100% acetonitrile) to 75% A and 25% B, a 10 min gradient to 5% A and 95% B, a 3 min isocratic under 5% A and 95% B. HR-ESI-MS analysis was conducted under negative scan mode over a mass range of 100 to 2000 Da. In the second, LC-MS was performed using an Agilent 1260 II Prime LC System with a Zorbax Eclipse XDB C18 column (100 mm × 2.1 mm × 3.5 µm, respectively), coupled to a 6475 triple quadrupole MS system. Metabolites were separated at a flow rate of 0.6 mL/min using a 2 min solvent gradient, from 70% solvent A (water with 0.02% acetic acid, 2 mM ammonium acetate, 0.1 mM ammonium fluoride) and 30% solvent B (100% methanol) to 40% A and 60% B, a 10 min gradient to 20% A and 80% B, and 1 min final gradient to 2% A and 98% B. Electrospray (jet stream) was conducted under negative scan mode over a mass range of 350 to 850 Da, followed by product ion and MRM modes for *m/z* 819.3, *m/z* 559.1, and *m/z* 494.9 using settings of 100 V fragmentor and 30 V collision energy. The masses represented saquayamycin A/B and two unique product ions (qualifiers), respectively.

### Transcriptional reporter systems

A promoter-less superfolder green fluorescence protein-encoding gene (with a terminator sequence upstream of the promoter cloning site to prevent read-through transcription) was amplified from pMC280 using primers pMS82MCS-F/pMS82MCS-R with a 60-64°C gradient annealing temperature. The resulting amplicons were cloned into the unique *EcoR*V site of the integrative plasmid pMS82 (57), yielding pMC281. The promoter region of the transcriptional activator *sqnR* (∼0.6 kb) was amplified using primers PsqnR-up1/PsqnR-up2 (62-68°C annealing temperature) and was cloned in the *Hind*III site upstream of the promoter-less green fluorescence protein-encoding gene. The resulting construct was verified by restriction analysis and sequencing using the universal M13-Reverse primer and was designated pMC282. This *sqnR* transcriptional reporter plasmid was then introduced into the WAC07094 Lsr2 knockdown strain where it integrated into the phiBT1 site in the chromosome.

We created an analogous transcriptional reporter construct for the *mtpA* gene. For this, we excised the *sqnR* promoter sequence from pMC282, and replaced it with the *mtpA* promoter. The *mtpA* promoter (∼0.2 kb) was amplified using primers PmtpA-HindIII/PmtpA-MfeI with a gradient annealing temperature from 66 to 70°C. The resulting reporter construct (pMC283) was confirmed by restriction analysis and sequenced using primer PmtpA-MfeI, before being introduced into WAC07094 strains. A positive control reporter strain for transcriptional activity was created by cloning the *ermE\** promoter upstream from the promoterless *gfp* gene and introducing this construct into different WAC07094 strains.

To visualize fluorescence for the different transcriptional reporter strains, the strains were inoculated onto solid growth medium and grown for 3 days before the colonies were imaged using a laser scanner (Typhoon™ FLA 9500) with an EGFP filter setting (473 nm). The resulting images were recorded without any further image processing.

### Creating deletion and complementation strains for *mtpAphoURP*

To simultaneously delete the *mtpA* and *phoURP* genes, DNA fragments flanking the four genes were amplified using primers pho-upKpnI/mtpA-dHindIII and pho-downSpeI/pho-downNotI with a 66-72°C gradient annealing temperature. A 3.1 kb-fragment (upstream of the four genes) was cloned into pIJ2925, whereas a 3.3 kb-fragment (downstream of the four genes) was cloned together with an apramycin resistance cassette (excised from pIJ773 using *Hind*III/*Spe*I) in pIJ777 (58). The downstream-flanking sequence and the apramycin resistance cassette were collectively excised using *Hind*III-*Psi*I and were introduced adjacent to the upstream flanking sequence at the *Hind*III-*Ssp*I sites. After construct integrity was confirmed by sequencing and digestion, the resulting plasmid (pMC284) was conjugated into WAC07094, and apramycin resistant exconjugants were selected for. Exconjugants were screened for double crossover events by PCR using primers phoP-INchk/phoR-INchk (where no product was expected in the event of a successful deletion; wild type chromosomal DNA was used as a positive control for these PCR checks) and oriT-2/mtpA-OUTchk with 65°C and 58°C annealing temperatures, respectively.

To dissect out the relative contribution of the four genes to the phenotype of the deletion mutant, four complementation constructs were created: (I) *mtpAphoURP,* using primers EXmtpA-PacI/pho-downKpnI, (II) *mtpAphoU,* using primers EXmtpA-PacI/PphoR-downKpnI, (III) *mtpA*, using primers EXmtpA-PacI/phoU-downKpnI, (IV) *phoURP*, using primers PhoU-downPacI/ pho-downKpnI. These amplicons were cloned into pIJ10257 (59) at the *Pac*I-*Kpn*I site (P*ermE*\* was excised out during cloning). The resulting constructs (pMC285-288, respectively) were introduced into the *mtpAphoURP* mutant by conjugation.

### Generating gene overexpression strains for select pleiotropic regulators

To individually overexpress *glnR* and *mtrA*, their genes were amplified from the WAC07094 genome using primers EXglnR-NdeI/EXglnR-XhoI or EXmtr-NdeI/EXmtr-PacI and a 58-65°C gradient annealing temperature. The resulting products were cloned into the *Nde*I/*Xho*I sites (*glnR*) or *Nde*I/*Pac*I site (*mtrA*) of the integrating plasmid pIJ10257, giving pMC289 and pMC290. Sequence integrity was confirmed by sequencing, after which the constructs were mobilized into both WAC07094 wild type and Lsr2 knockdown strains through conjugation. Overexpressing the constitutive *afsQ1*D52E was achieved using the thiostrepton-inducible pIJ6902-based construct (23).

### Assessing saquayamycin production during liquid culturing

Dilutions of WAC07094 or CCESR44 spores (10^7^-10^3^ or 10^7^-10^2^ viable spores/mL, respectively) were inoculated into 5 mL of Bennett’s liquid medium in 28 mL universal bottles. The cultures were then shaken (200 RPM) at 30°C for 4 days. Metabolites were extracted from the cultures using equal volumes of ethyl acetate. The solvent was evaporated *in vacuo* and the metabolites were resuspended in 50 µL DMSO. For the bioassays, 10 µL of metabolite extract were applied to the filter discs, as described above.

### Genetic manipulation of strain 3212.3 for Lsr2 activity modulation and saquayamycin biosynthetic gene disruption

To modulate Lsr2 activity in CCESR44, the Lsr2 knockdown construct was introduced via intergeneric conjugation. Approximately 10^8^ viable spores were mixed with 1.5 mL of overnight cultures of *E. coli* ET12567/pUZ8002, before being spread on ISP4 agar supplemented with 20 mM MgCl2. After overnight growth, plates were overlaid with apramycin to select for exconjugants.

To disrupt saquayamycin biosynthesis in strain 3212.3, we took advantage of the fact that the *sqnHI* sequence was 98% identical (over 2.3 kb) between WAC07094 and CCESR44. We introduced the pIJ10700+*sqnHI* plasmid (pMC341) into CCESR44 through conjugation, selecting for hygromycin resistant exconjugants.

## ACKNOWLEDGMENTS

We would like to thank the Wright lab for their gift of WAC07094 and sharing its Illumina-based genome sequence, Dr. Linda Kinkel for providing access to *Streptomyces* isolate CCESR44, Dan Browne for his generous assistance with genome assembly, Dr. Brian Golding for assistance with computing access, and John Fast and the Kidd lab for technical assistance with the LC-triple quadrupole mass spectrometry. The sequencing of *Streptomyces* isolate CCESR44 (previously annotated as *Streptomyces* sp. 3212.3) was conducted by the U.S. Department of Energy Joint Genome Institute (https://ror.org/04xm1d337) (proposal: 10.46936/10.25585/60001044), a DOE Office of Science User Facility that is supported by the Office of Science of the U.S. Department of Energy operated under Contract No. DE-AC02-05CH11231. This work was supported by a PacBio SMRT sequencing grant to H, and by a CIHR Project grant (162340) to MAE.

## REFERENCES

1. Ramírez-Rendon D, Passari AK, Ruiz-Villafán B, Rodríguez-Sanoja R, Sánchez S, Demain AL. 2022. Impact of novel microbial secondary metabolites on the pharma industry. Appl Microbiol Biotechnol 106:1855–1878.

2. Newman DJ, Cragg GM. 2020. Natural products as sources of new drugs over the nearly four decades from 01/1981 to 09/2019. J Nat Prod 83:770–803.

3. Walsh CT. 2018. Nature builds macrocycles and heterocycles into its antimicrobial frameworks: Deciphering Biosynthetic Strategy. ACS Infect Dis 4:1283–1299.

4. Chater KF. 2006. *Streptomyces* inside-out: a new perspective on the bacteria that provide us with antibiotics. Philos Trans R Soc Lond B Biol Sci 361:761–8.

5. Urem M, Świątek-Połatyńska MA, Rigali S, van Wezel GP. 2016. Intertwining nutrient-sensory networks and the control of antibiotic production in *Streptomyces*. Molecular Microbiology 102:183–195.

6. McCormick JR, Flärdh K. 2012. Signals and regulators that govern *Streptomyces* development. FEMS Microbiol Rev 36:206–31.

7. Rigali S, Titgemeyer F, Barends S, Mulder S, Thomae AW, Hopwood DA, van Wezel GP. 2008. Feast or famine: the global regulator DasR links nutrient stress to antibiotic production by *Streptomyces*. EMBO Rep 9:670–5.

8. Horinouchi S, Beppu T. 2007. Hormonal control by A-factor of morphological development and secondary metabolism in *Streptomyces*. Proc Jpn Acad Ser B Phys Biol Sci 83:277–95.

9. Martín JF, Liras P. 2020. The balance metabolism safety net: integration of stress signals by interacting transcriptional factors in *Streptomyces* and related actinobacteria. Front Microbiol 10:3120.

10. Reuther J, Wohlleben W. 2007. Nitrogen metabolism in *Streptomyces coelicolor*: transcriptional and post-translational regulation. J Mol Microbiol Biotechnol 12:139–46.

11. Martín JF, Liras P. 2021. Molecular mechanisms of phosphate sensing, transport and signalling in *Streptomyces* and related actinobacteria. Int J Mol Sci 22:1129.

12. Gehrke EJ, Zhang X, Pimentel-Elardo SM, Johnson AR, Rees CA, Jones SE, Hindra, Gehrke SS, Turvey S, Boursalie S, Hill JE, Carlson EE, Nodwell JR, Elliot MA. 2019. Silencing cryptic specialized metabolism in *Streptomyces* by the nucleoid-associated protein Lsr2. eLife 8: e47691.

13. Santos-Beneit F, Rodríguez-García A, Sola-Landa A, Martín JF. 2009. Cross-talk between two global regulators in *Streptomyces*: PhoP and AfsR interact in the control of *afsS*, *pstS* and *phoRP* transcription. Mol Microbiol 72:53–68.

14. Liu G, Chater KF, Chandra G, Niu G, Tan H. 2013. Molecular regulation of antibiotic biosynthesis in *Streptomyces*. Microbiol Mol Biol Rev 77:112–43.

15. Gordon BRG, Imperial R, Wang L, Navarre WW, Liu J. 2008. Lsr2 of *Mycobacterium* represents a novel class of H-NS-like proteins. J Bacteriol 190:7052–7059.

16. Uchida T, Imoto M, Watanabe Y, Miura K, Dobashi T, Matsuda N, Sawa T, Naganawa H, Hamada M, Takeuchi T. 1985. Saquayamycins, new aquayamycin-group antibiotics. J Antibiot (Tokyo) 38:1171–81.

17. Erb A, Luzhetskyy A, Hardter U, Bechthold A. 2009. Cloning and sequencing of the biosynthetic gene cluster for saquayamycin Z and galtamycin B and the elucidation of the assembly of their saccharide chains. Chembiochem 10:1392–401.

18. Blin K, Shaw S, Steinke K, Villebro R, Ziemert N, Lee SY, Medema MH, Weber T. 2019. antiSMASH 5.0: updates to the secondary metabolite genome mining pipeline. Nucleic Acids Res 47:W81–W87.

19. Salem SM, Weidenbach S, Rohr J. 2017. Two cooperative glycosyltransferases are responsible for the sugar diversity of saquayamycins Isolated from *Streptomyces* sp. KY 40-1. ACS Chem Biol 12:2529–2534.

20. Ghorbel S, Kormanec J, Artus A, Virolle MJ. 2006. Transcriptional studies and regulatory interactions between the *phoR-phoP* operon and the *phoU*, *mtpA*, and *ppk* genes of *Streptomyces lividans* TK24. J Bacteriol 188:677–86.

21. Sola-Landa A, Rodríguez-García A, Franco-Domínguez E, Martín JF. 2005. Binding of PhoP to promoters of phosphate-regulated genes in *Streptomyces coelicolor*: identification of PHO boxes. Mol Microbiol 56:1373–85.

22. Sola-Landa A, Rodríguez-García A, Apel AK, Martín JF. 2008. Target genes and structure of the direct repeats in the DNA-binding sequences of the response regulator PhoP in *Streptomyces coelicolor*. Nucleic Acids Res 36:1358–68.

23. Daniel-Ivad M, Hameed N, Tan S, Dhanjal R, Socko D, Pak P, Gverzdys T, Elliot MA, Nodwell JR. 2017. An engineered allele of *afsQ1* facilitates the discovery and investigation of cryptic natural products. ACS Chem Biol 12:628–634.

24. Tatusova T. 2016. Update on genomic databases and resources at the National Center for Biotechnology Information. Methods Mol Biol 1415:3–30.

25. Hadjithomas M, Chen IA, Chu K, Huang J, Ratner A, Palaniappan K, Andersen E, Markowitz V, Kyrpides NC, Ivanova NN. 2017. IMG-ABC: new features for bacterial secondary metabolism analysis and targeted biosynthetic gene cluster discovery in thousands of microbial genomes. Nucleic Acids Res 45:D560–D565.

26. Alanjary M, Steinke K, Ziemert N. 2019. AutoMLST: an automated web server for generating multi-locus species trees highlighting natural product potential. Nucleic Acids Res 47:W276–W282.

27. Price-Whelan A, Dietrich LE, Newman DK. 2006. Rethinking ’secondary’ metabolism: physiological roles for phenazine antibiotics. Nat Chem Biol 2:1–8.

28. Eberl L. 1999. N-acyl homoserinelactone-mediated gene regulation in gram-negative bacteria. Syst Appl Microbiol 22:493–506.

29. Sánchez L, Braña AF. 1996. Cell density influences antibiotic biosynthesis in *Streptomyces clavuligerus*. Microbiology (Reading) 142:1209–1220.

30. Biarnes-Carrera M, Breitling R, Takano E. 2015. Butyrolactone signalling circuits for synthetic biology. Curr Opin Chem Biol 28:91–8.

31. Vashishtha AK, Wang J, Konigsberg WH. 2016. Different divalent cations alter the kinetics and fidelity of DNA polymerases. J Biol Chem 291:20869–20875.

32. Cromie MJ, Shi Y, Latifi T, Groisman EA. 2006. An RNA sensor for intracellular Mg(2+). Cell 125:71–84.

33. Dann CE, 3rd, Wakeman CA, Sieling CL, Baker SC, Irnov I, Winkler WC. 2007. Structure and mechanism of a metal-sensing regulatory RNA. Cell 130:878–92.

34. Forrest D. 2019. Unusual relatives of the multisubunit RNA polymerase. Biochem Soc Trans 47:219–228.

35. Walker TE, Shirzadeh M, Sun HM, McCabe JW, Roth A, Moghadamchargari Z, Clemmer DE, Laganowsky A, Rye H, Russell DH. 2022. Temperature regulates stability, ligand binding (Mg(2+) and ATP), and stoichiometry of GroEL-GroES complexes. J Am Chem Soc 144:2667–2678.

36. Will WR, Whitham PJ, Reid PJ, Fang FC. 2018. Modulation of H-NS transcriptional silencing by magnesium. Nucleic Acids Res 46:5717–5725.

37. Groisman EA, Chan C. 2021. Cellular adaptations to cytoplasmic Mg(2+) limitation. Annu Rev Microbiol 75:649–672.

38. Pontes MH, Groisman EA. 2018. Protein synthesis controls phosphate homeostasis. Genes Dev 32:79–92.

39. Lee J, Wu J, Deng Y, Wang J, Wang C, Wang J, Chang C, Dong Y, Williams P, Zhang L-H. 2013. A cell-cell communication signal integrates quorum sensing and stress response. Nature Chemical Biology 9:339–343.

40. Zhang X, Andres SN, Elliot MA. 2021. Interplay between nucleoid-associated proteins and transcription factors in controlling specialized metabolism in *Streptomyces*. mBio 12:10.1128/mbio.01077-21.

41. Kharel MK, Pahari P, Shepherd MD, Tibrewal N, Nybo SE, Shaaban KA, Rohr J. 2012. Angucyclines: Biosynthesis, mode-of-action, new natural products, and synthesis. Nat Prod Rep 29:264–325.

42. Caraballo-Rodríguez AM, Dorrestein PC, Pupo MT. 2017. Molecular inter-kingdom interactions of endophytes isolated from *Lychnophora ericoides*. Sci Rep 7:5373.

43. Schneemann I, Kajahn I, Ohlendorf B, Zinecker H, Erhard A, Nagel K, Wiese J, Imhoff JF. 2010. Mayamycin, a cytotoxic polyketide from a *Streptomyces* strain isolated from the marine sponge *Halichondria panicea*. J Nat Prod 73:1309–12.

44. Fujikawa H, Akimoto R. 2011. New blue pigment produced by *Pantoea agglomerans* and its production characteristics at various temperatures. Appl Environ Microbiol 77:172–8.

45. Sambrook J, Russell DW. 2001. Molecular cloning : a laboratory manual, 3rd ed. Cold Spring Harbor Laboratory Press, Cold Spring Harbor, N.Y.

46. MacNeil DJ, Gewain KM, Ruby CL, Dezeny G, Gibbons PH, MacNeil T. 1992. Analysis of *Streptomyces avermitilis* genes required for avermectin biosynthesis utilizing a novel integration vector. Gene 111:61–8.

47. Kieser T, Bibb MJ, Buttner MJ, Chater KF, Hopwood DA. 2000. Practical Streptomyces genetics. John Innes Foundation, Norwich.

48. Shirling EB, Gottlieb D. 1966. Methods for characterization of *Streptomyces* species. Int J Syst Bacteriol 16:313–340.

49. Waglechner N, McArthur AG, Wright GD. 2019. Phylogenetic reconciliation reveals the natural history of glycopeptide antibiotic biosynthesis and resistance. Nat Microbiol 4:1862–1871.

50. Kolmogorov M, Yuan J, Lin Y, Pevzner PA. 2019. Assembly of long, error-prone reads using repeat graphs. Nat Biotechnol 37:540–546.

51. Li H. 2018. Minimap2: pairwise alignment for nucleotide sequences. Bioinformatics 34:3094–3100.

52. Vaser R, Sović I, Nagarajan N, Šikić M. 2017. Fast and accurate de novo genome assembly from long uncorrected reads. Genome Res 27:737–746.

53. Schneider CA, Rasband WS, Eliceiri KW. 2012. NIH Image to ImageJ: 25 years of image analysis. Nat Methods 9:671–5.

54. Jiang H, Lei R, Ding SW, Zhu S. 2014. Skewer: a fast and accurate adapter trimmer for next-generation sequencing paired-end reads. BMC Bioinformatics 15:182.

55. Langmead B, Trapnell C, Pop M, Salzberg SL. 2009. Ultrafast and memory-efficient alignment of short DNA sequences to the human genome. Genome Biol 10:R25.

56. Robinson JT, Thorvaldsdóttir H, Winckler W, Guttman M, Lander ES, Getz G, Mesirov JP. 2011. Integrative genomics viewer. Nat Biotechnol 29:24–6.

57. Gregory MA, Till R, Smith MC. 2003. Integration site for *Streptomyces* phage phiBT1 and development of site-specific integrating vectors. J Bacteriol 185:5320–3.

58. Gust B, Challis GL, Fowler K, Kieser T, Chater KF. 2003. PCR-targeted *Streptomyces* gene replacement identifies a protein domain needed for biosynthesis of the sesquiterpene soil odor geosmin. Proc Natl Acad Sci U S A 100:1541–6.

59. Hong HJ, Hutchings MI, Hill LM, Buttner MJ. 2005. The role of the novel Fem protein VanK in vancomycin resistance in *Streptomyces coelicolor*. J Biol Chem 280:13055–61.

